# Four different mechanisms for switching cell polarity

**DOI:** 10.1101/2020.06.16.155952

**Authors:** Filipe Tostevin, Manon Wigbers, Lotte Søgaard-Andersen, Ulrich Gerland

**Affiliations:** Physics of Complex Biosystems, Physics Department, Technical University of Munich, James-Franck-Str. 1, 85748 Garching, Germany; Arnold Sommerfeld Center for Theoretical Physics and Center for Nanoscience, Ludwig-Maximilians-Universität München, Theresienstr. 37, 80333 München, Germany; Max Planck Institute for Terrestrial Microbiology, Karl-von-Frisch-Str. 10, 35043 Marburg, Germany

**Author notes:** Contributed equally to this work (Filipe Tostevin, Manon Wigbers).

**Keywords:** cell polarity, signal transduction, toggle switch

## Abstract

The mechanisms and design principles of regulatory systems establishing stable polarized protein patterns within cells are well studied. However, cells can also dynamically control their cell polarity. Here, we ask how an upstream signaling system can switch the orientation of a polarized pattern. We use a mathematical model of a core polarity system based on three proteins as the basis to study different mechanisms of signal-induced polarity switching. The analysis of this model reveals four general classes of switching mechanisms with qualitatively distinct behaviors: the transient oscillator switch, the reset switch, the prime-release switch, and the push switch. Each of these regulatory mechanisms effectively implements the function of a spatial toggle switch, however with different characteristics in their nonlinear and stochastic dynamics. We identify these characteristics and also discuss experimental signatures of each type of switching mechanism.

---

Cell polarity is manifested in molecular and morphological asymmetries of the cell. From bacterial to mammalian cells, cell polarity is essential in a multitude of functional contexts, including cell migration, cell division and differentiation, cell-cell signaling, development and tissue homeostasis [1, 2]. One fundamental question related to cell polarity is how an initially symmetrical cell can establish a polarized state and subsequently maintain it [3]. However, cells are also known to dynamically change their polarity, e.g. reversing polarity in response to external or internal signals to control motility [4–6]. This raises a second fundamental question: Which mechanisms permit reliable switching of cell polarity?

The first question, about establishing and maintaining cell polarity, is well studied, both on the conceptual level with theoretical approaches and on the experimental level by characterizing model systems. The polarization of an initially nonpolarized cell is a symmetry breaking phenomenon: In the case of essentially isotropic cells, e.g. budding yeast or epithelial cells [3], the continuous angular symmetry is broken by polarization, whereas discrete symmetry breaking occurs for rod-shaped bacterial cells [7]. Symmetry breaking can occur spontaneously [8], but is often controlled by upstream guiding cues [9]. While the detailed molecular mechanisms underlying cell polarization differ between organisms, they often incorporate conserved G-protein based signaling systems that use multiple feedback interactions to generate asymmetric distributions on the cell membrane via a Turing instability [10]. A class of simple networks that can achieve cell polarization was explored in a synthetic biology study [11], which first showed computationally that all such networks feature one or more of the three minimal motifs ‘positive feedback’, ‘mutual inhibition’, or ‘inhibition with positive feedback’, and that combinations of these motifs generally polarize more reliably. The study also corroborated the latter finding experimentally, recapitulating the basic principles underlying the establishment and maintenance of cell polarization in engineered systems. Taken together, these and other results address many aspects of the first question raised above. By comparison, significantly less is known about the second question on the dynamical control of cell polarity.

Dynamically changing cell polarity is widely observed and studied in eukaryotic model systems such as migrating neutrophils [12] and amoebae [5], as well as melanoma cells [13]. Depending on the system, its genetic makeup, and the environment, cells display a variety of dynamical patterns. For instance, melanoma cells either randomly polarize into frequently changing directions, or reverse cell polarity in an oscillatory fashion, or they persistently maintain cell polarity [13]. The dynamical control of cell polarity involves signaling. For instance, cell polarity changes can be coupled to internal signals, as in the case of yeast, where the dynamics of cell polarity is co-regulated by the cell cycle [14]. Often, cells get a directional cue from the environment governing the direction of their response [5]. However, cells can also respond to non-directional cues. For instance, a temporally decreasing chemoattractant signal triggers reversals of cell polarity in neutrophils, even in the absence of a spatial concentration gradient [12]. Which mechanisms permit such reversals induced by a non-directional signal?

Rod-shaped bacteria display much of the eukaryotic phenomenology and serve as paradigmatic model systems. For instance, the Min system, used by *Escherichia coli* to localize the septum prior to cell division [15], constitutes a prime example of autonomous cell polarity oscillations. Its underlying molecular network, based on three Min proteins, was successfully reconstituted *in vitro* [16]. On a conceptual level, the cell polarity oscillations of the Min system are analogous to those of the melanoma cells, also with respect to the basic regulatory scheme, whereby a bistable system can be turned into an oscillator via slow negative feedback [13, 17]. For signal-induced (rather than autonomous) polarity reversal, the Mgl/Rom system of *Myxococcus xanthus* constitutes a prime example. Here the cell polarity, marked by MglA, undergoes intermittent reversals triggered by the upstream Frz signaling system [6, 18]. The cell polarity reversals are accompanied by reversals in the direction of cell motion, enabling motility patterns that are crucial for predatory behavior and fruiting body formation [19, 20].

Recently, Guzzo et al [21] identified the response regulator FrzX as a mediator of the Frz reversal signal to the Mgl system, and proposed a mechanism for how FrzX can interact with the three core polarity proteins to trigger polarity reversals. Here, we take this study as a starting point to explore the question of signal-induced polarity switching on a more general level. Rather than focusing on one particular mechanism, we aim to identify the distinct classes of switching mechanisms and their underlying working principles. We find four distinct classes of mechanisms that can occur for different signaling regimes. We demonstrate that some are sensitive to the amplitude and duration of the input signal but relatively robust to intrinsic molecular noise, while others are less sensitive to signal variability but more susceptible to noise. These and other features allow us to identify experimental signatures that can be used to discriminate between the four classes of mechanisms *in vivo*.

## RESULTS

We consider a cell polarity defined by an asymmetric distribution of a certain ‘polarity marker’ *A*. The polarity marker has the regulatory role to direct the spatial localization or activity of downstream processes. For instance, MglA in *M. xanthus* is a polarity marker that localizes at one of the cell poles and activates the motility machinery to determine the direction of cell motion [18]. Similarly, Cdc42 is a polarity marker in yeast and other eukaryotic cells [22]. A module consisting of the polarity marker and other regulatory proteins has the ability to establish and maintain a polarized distribution of *A*. This module, which we refer to as the ‘core polarity system’, receives input from a signaling pathway via a signaling protein *X*. We stipulate that the ‘full polarity system’ consisting of *X* and the core system can implement the function of signal-induced polarity switching (Fig. 1).

**FIG. 1:**
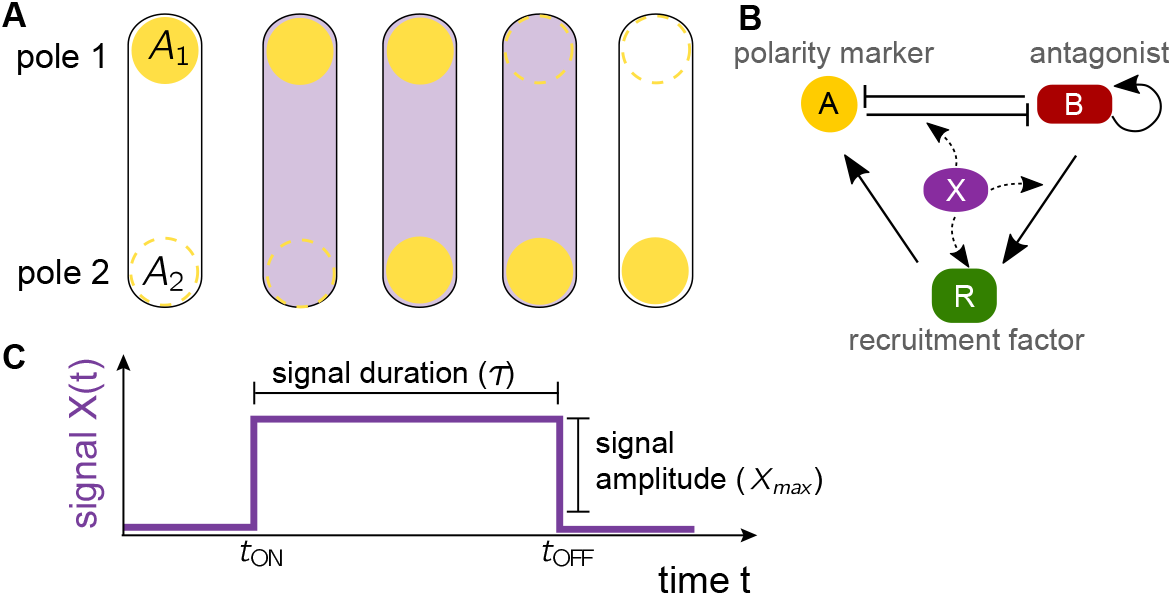
Signal-induced polarity switching. **A** Schematic representation of a rod-shaped cell with polarity marker *A* shown in yellow. Proteins can either be bound to the poles or diffuse in the cytoplasm. The abundances of the polarity marker at the two poles are denoted by *A*_1_ and *A*_2_. The release of a signal protein *X* in the cytoplasm, shown in purple, can lead to a polarity reversal, such that the polarity marker switches from pole 1 to pole 2. **B** Schematic representation of the molecular interactions of the polarity model. The polarity marker *A* and its antagonist *B* inhibit each others binding to the pole. *B* can cooperatively recruit itself to the pole and promotes binding of the recruitment factor *R*, which in turn recruits *A*. Dashed lines indicate exemplary hypothetical interactions of the signal protein *X* with the polarity proteins. **C** The switching signal is implemented as a pulse in the total amount of *X*, parameterized by the signal duration *τ* and signal amplitude *X*_max_.

To explore mechanisms for signal-induced polarity switching, we consider a symmetric cell with a polarity marker that localizes only at its two cell poles ‘1’ and ‘2’ (Fig. 1A), while it rapidly diffuses in the cytoplasm. This simplest scenario is a good approximation for the *M. xanthus* polarity system [21] and suffices to reveal general principles of signal-induced polarity switching, as we will see below. The distribution of *A* is then characterized by quantifying its abundance at pole ‘1’ and ‘2’, as well as in the cytoplasm, and the time-dependent cell polarity can be defined as

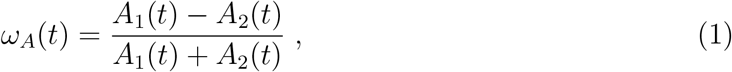

where *A*_1_(*t*) and *A*_2_(*t*) are the time-dependent abundances of *A* at the poles. Hence, *ω_A_* > 0 corresponds to a higher abundance of *A* at pole 1 than at pole 2, and vice versa for *ω_A_* < 0, such that a reversal of cell polarity is marked by a change of sign in *ω_A_*(*t*).

### Model for a switchable polarity system

To obtain our working model, we generalize the recently proposed model of the *M. xanthus* polarity system [21]. This model involves the ‘antagonist’ *B* to the polarity marker *A*, as well as a third protein species, the ‘recruitment factor’ *R* (representing MglB and RomR, respectively). The network of interactions between *A*, *B*, and *R* is shown in Fig. 1B. Besides the mutual inhibition between *A* and *B*, it involves self-recruitment of *B*, as well as indirect recruitment of *A* by *B* via *R*. The full dynamics of the interactions between *A*, *B*, and *R* at the poles is described by [21]

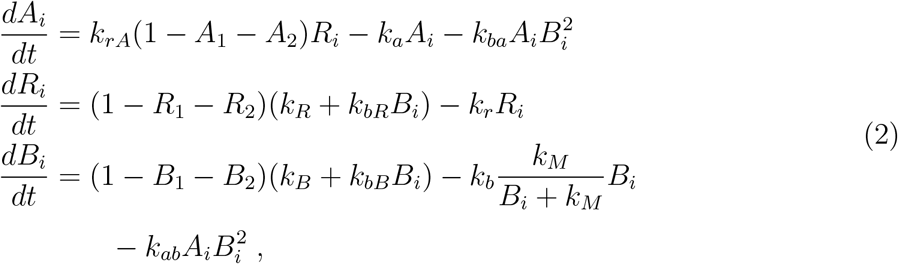

using the same convention for *B* and *R* as for *A*, i.e. *B_i_* and *R_i_* denote the abundances at the poles (*i* = 1,2). Eq. 2 assumes that the total abundances of *A*, *B*, and *R* in the cell are approximately constant, at least on the relevant timescale of polarity reversals. These total values are set to one by choosing appropriate units for the abundances. The dynamics in the cytoplasm is then obtained from the dynamics of the polar abundances, e.g. the cytoplasmic abundance of *A* is 1 − *A*_1_ − *A*_2_. In total, the interactions between *A*, *B*, and *R* are specified by 10 rate constants and one saturation parameter. *R* binds to the cell poles with rate *k_R_* where it locally recruits *A* with rate *k_rA_*. *B* binds at the intrinsic rate *k_B_* to the poles, where it recruits both itself, at rate *k_bB_*, and *R* at rate *k_bR_*. At the same time, *A* can displace *B* from the pole and vice versa with a rate *k_ab_* and *k_ba_*, respectively. All three proteins can also spontaneously unbind from the poles, with the corresponding rates *k_a_*, *k_r_*, and *k_b_*, but the unbinding of *B* is slowed in presence of more *B* (with the saturation parameter *k_M_* determining the characteristic abundance for this feedback effect).

The positive feedback from *B* onto its own localization together with the mutual inhibition of *A* and *B* allow this model to spontaneously generate a stable asymmetry in the protein abundances at the two poles. Polarity schemes based on mutual antagonism also play a role in polarity establishment of the PAR system [23] determining the anterior-posterior axis in *C. elegans*, and the Rac-Rho system regulating front-rear polarity in mammalian cells [13]. Here, we use Eq. 2 to describe the deterministic dynamics of the core polarity system. To explore noise effects due to the relatively low copy numbers of regulatory proteins within cells, we also devised a stochastic model based on stochastic differential equations, see ‘Materials and Methods’. The noise strength in this model is determined by an effective “copy number” parameter *N*, with *N* → ∞ recovering the deterministic dynamics and noise strength increasing with decreasing *N*.

The signaling protein *X* mediates a non-directional signal that interacts with the core polarity system (Fig. 1B) to induce polarity switching (in *M. xanthus*, *X* corresponds to phosphorylated FrzX [21]). We assume the total amount *X_t_* of *X* to have a step-like pulse form (Fig. 1C), parameterized by an amplitude *X*_max_ and duration *τ*. While step-like pulses are a reasonable assumption, given that signals change via (rapid) protein modifications rather than (slow) changes in protein levels, we will also study the effect of more gradual changes further below. In the model of [21], cytoplasmic *X* is recruited to the poles by the antagonist *B* with rate *k_X_* and spontaneously unbinds with rate *k_x_*, such that its polar abundances change according to

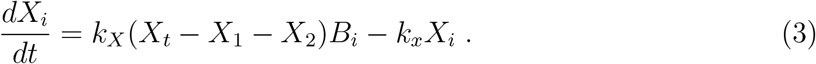

In order to systematically explore the possible mechanisms by which polar *X* may interact with the core polarity system, we allowed *X* to regulate each one of the 11 parameters in Eq. 2. We allowed for both positive and negative regulation, thus obtaining 22 different candidate models for a switchable polarity system. In each case, one parameter *k_j_* depends on *X_i_* while the others are not affected. For a positive regulation, we have

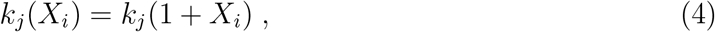

and for a negative

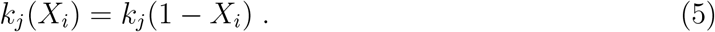

Hence, a candidate signaling scenario is parameterized by (i) which parameter *k_j_* is regulated by *X*, (ii) whether the regulation is enhancing or repressive, and (iii) the amplitude and duration of the pulse, as illustrated in Fig. 2A.

**FIG. 2:**
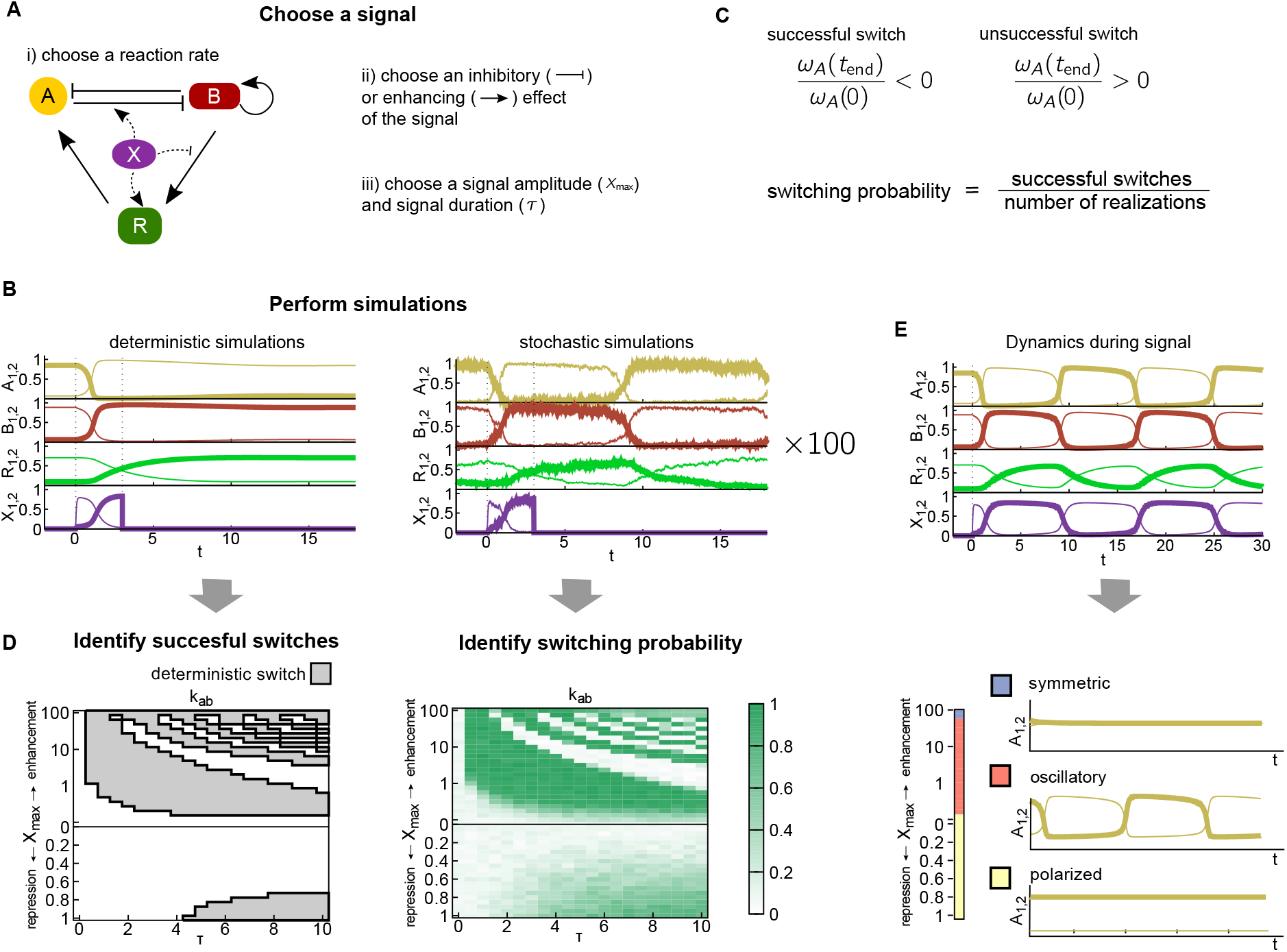
Schematic representation of the workflow. **A** Switching signals are parameterized by the choice of i) a reaction rate it acts on, ii) an inhibitory or enhancing effect and iii) the amplitude *X*_max_ and duration *τ* of the transient signal. *X* can act on any of the 11 parameters of the polarity model. **B** Example of a deterministic and stochastic simulation before, during and after the signal. The signal is applied between *t* = 0 and *t* = 3. Thick lines indicate the concentrations of *A* (yellow), *B* (red), *R* (green) and *X* (purple) at pole 1, and thin lines at pole 2. **C** Switching is evaluated by comparing the signs of the asymmetry in *A* before *ω*(0) and after the switch *ω*(*t*_END_). For the stochastic simulation a switching probability is calculated from 100 trajectories. **D** Switching regimes are plotted in phase space as a function of *X*_max_ and *τ* for the modification of each model parameter. For the deterministic model, successful switches are shown by the gray regions with a black outline, for the stochastic model switching probabilities are shown in green. **E** The state of the system during the signal is identified by simulating the deterministic model with the signal applied for the duration of the simulations. The dynamics is classified into three states: symmetric (blue), oscillatory (orange) and polarized (yellow).

### Identifying functional switching scenarios

To test a candidate signaling scenario for its ability to induce polarity switching, we simulate the dynamics (both deterministic and stochastic) of the model. The output of a simulation is a set of time-dependent abundances of the four proteins *A*, *B*, *R*, and *X* at the two poles (Fig. 2B). Each simulation run has three phases. First, we simulate the polarity model, Eq. 2, in the absence of signaling input (*X_t_* = 0). In this condition, the system reaches a stable polarized configuration. At *t* = 0, we then switch to *X_t_* = *X*_max_ for a duration *τ*, after which the simulation is continued with *X_t_* set to zero again. We then compare the polarization of the cell at the time when the signal is initiated (*t* = 0) with a time point after the removal of the signal (*t*_end_ = 30 was chosen to allow for the system to fully relax back to a polarized steady state). The candidate signaling scenario is considered to generate a successful switch if the signs of *ω_A_*(0) and *ω_A_*(*t*_end_) were different (i.e., the initial and final polarity states were different), and unsuccessful otherwise (Fig. 2C). For the stochastic dynamics, we estimated the switching probability from 100 simulation runs (Fig. 2B,C). We repeated this procedure for each signaling scenario with a range of *X*_max_ and *τ* values, generating deterministic and stochastic phase diagrams delineating the functional regimes in (*τ, X*_max_)-space (Fig. 2D).

### Characterizing succesful switching scenarios

Fig. 3 shows the resulting phase diagrams, each representing regulation via one of the 11 model parameters and including both enhancing and repressive regulatory effects. Here, the deterministic regimes of successful polarity reversals (solid black lines) are superimposed with the stochastic switching probabilities (green shading). We identified at least one range of signal parameters with successful polarity reversals in each of the phase diagrams. That is, it is possible for *X* to induce reversals by regulating any of the interactions of the polarity proteins, provided that the profile of the signal pulse *X_t_* is chosen appropriately. Surprisingly, in most cases reversals can be observed when *X* acts either positively *or* negatively. For example, reversals can be induced by *X* either enhancing or repressing the strength of *B* self-recruitment via the parameter *k_bB_*.

**FIG. 3:**
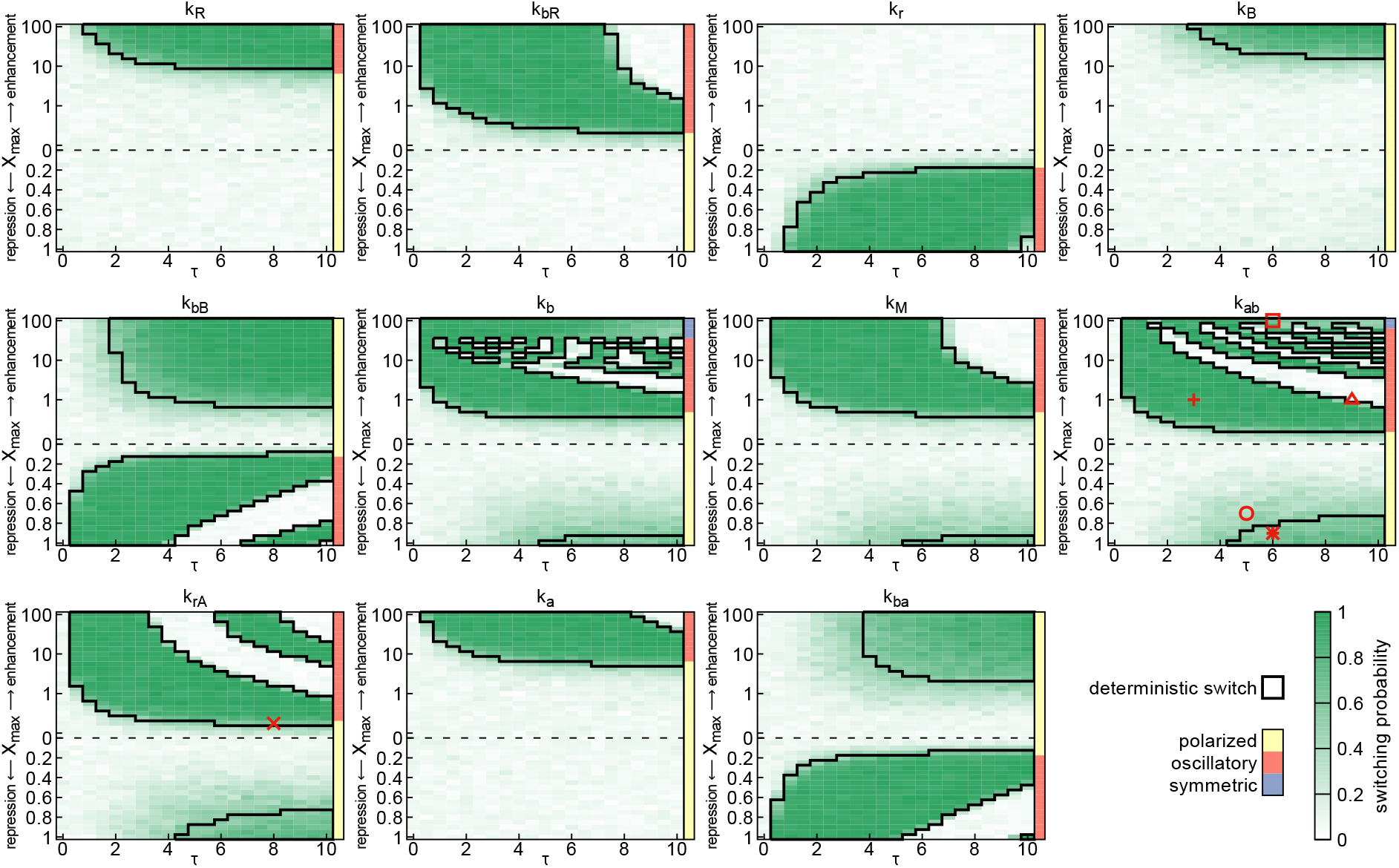
Switching regimes for each of the model parameters. Regions in which the deterministic model shows switches are indicated by thick black outlines. The green shading shows the switching probability of the stochastic model with *N* = 10^3.75^. The upper half of the phase diagram shows results for a signal that enhances the reaction rate, and the lower half for a repression of the rate. The colored bars to the right of each panel indicate the class of dynamics when the corresponding amplitude of signal is applied, with yellow for polarized, orange for oscillatory and blue for symmetric polar distribution of *A*. The red symbols indicate the signal amplitude and duration of the trajectories shown in Fig. 4

Polarity is highly sensitive to regulation of some parameters (e.g. *k_bB_*, *k_ba_*), with switching occurring for most signal profiles. These parameters tend to be those involved in the key interactions of Fig. 1B, including the nonlinear feedbacks in *B* recruitment, *A* recruitment by *R*, and *A*-*B* mutual antagonism, which together are crucial for the establishment and maintenance of polarity. For parameters that are more peripheral to the interaction network, in particular the spontaneous binding and dissociation rates (e.g. *k_R_*, *k_B_*), switching occurs only in small regions of high-amplitude signals.

Fig. 3 reveals two qualitatively different patterns in the signaling regimes generating switching: solid regions, in which switching is insensitive to *X*_max_ and *τ* provided these exceed a threshold; and alternating bands of successful and unsuccessful switching regions, in which the system remains sensitive to the values of *X*_max_ and *τ*. Intuitively, alternating bands would be expected to occur, if the system dynamics become oscillatory in presence of the signal, since Fig. 3 only compares the initial and final state, such that for instance it does not discriminate between trajectories in which polarity is never reversed, and those in which polarity reverses twice.

To investigate the switching mechanism in the succesful parameter regimes, we examined trajectories of the system for different signals. For a trajectory within a banded region (plus symbol in Fig. 3), we see that once the signal is applied, *A* rapidly relocates to the opposite pole, followed by *B* and on a slower timescale *R* (Fig. 4A, solid lines). If a signal with the same amplitude *X*_max_ is applied for a longer time (open triangle in Fig. 3), a second switch takes place (Fig. 4A, dashed lines). Hence the width of the bands is determined by the timescale of *R* reorientation. This particular case, where *X* enhances *k_ab_*, is precisely the relaxation oscillator dynamics reported in [21].

**FIG. 4:**
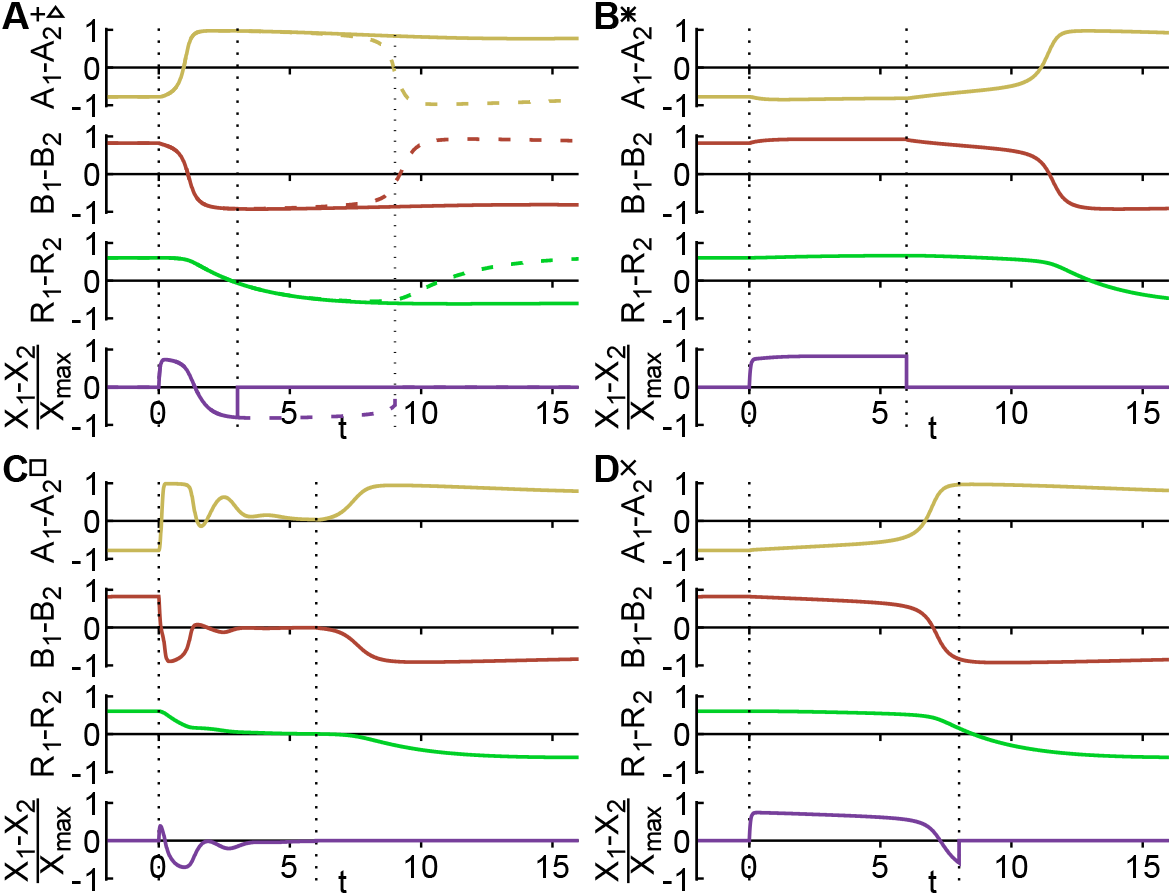
Trajectories of the model during switches, classified as four different switching classes. Signal parameters *X*_max_ and *τ* and the parameter modified are indicated by the corresponding symbols in Fig. 3. Vertical dashed lines indicate the period during which the signal is present. **A** Relaxation oscillator. For a short signal (plus-symbol), the polarity switches during the applied signal as shown by the solid lines. For a longer signal (open triangle), the system switches a second time as shown by the dashed lines. **B** Prime-release switch. During the signal the polarity is unchanged, but switches after the signal is released. **C** Reset switch. During the signal, the system relaxes to a symmetric distribution of the polarity marker and establishes a reversed polarity after the signal is removed. **D** Push switch. The system switches while the signal is applied and does not switch back when the signal is applied longer.

For a trajectory in the non-band signal regime (star symbol in Fig. 3), the system rapidly reaches a new steady state (with the same polarity) when the signal is applied (Fig. 4B). The polarity reversal occurs after, and appears to be initiated by, the removal of the signal. To confirm that there are no longer-period oscillations during the signal period, we examined the dynamics with a signal of the same amplitude for a long duration (*τ* = 100). The system remained stably polarized throughout this duration. Thus, this switching mechanism is qualitatively different from the relaxation oscillator reported previously. Switching is insensitive to the signal duration *τ*, provided that it is above a threshold value. We interpret this threshold as meaning that the signal must be present for long enough to prime the system to switch, and refer to this mechanism as a “prime-release” switch.

We then examined trajectories over the entire signal space and determined the order in which the polarity of *A*, *B* and *R* reversed, defined by the times at which their asymmetry *ω* becomes zero. For almost all regimes with reversals the same order was observed (Fig. S1): first *A*, then *B*, and finally *R*. This suggests that the underlying dynamics of the trajectory between the two polarity states is similar in different switching regimes. In some limited regimes, for particularly high-amplitude signals, reversal of first *B* and then *A* was observed. However, these reversals were almost simultaneous. In some regimes reversals of *A* and *B* but not *R* occurred. In these cases, the polarity oscillations of *A* and *B* were so fast that a second reversal was initiated before the much slower dynamics of *R* could catch up to the new polarity state.

### Classification of switching mechanisms

We next examined the dynamics during persistent signals for all regulations and signal amplitudes (Fig. 2E). We identified three classes of behavior (Fig. S2), reflecting qualitatively different topologies of the model’s state space as shown schematically in Fig. 5A. These are

i. static asymmetrically polarized protein distributions, corresponding to bistable state space with the two stable states representing the two possible orientations of polarization;
ii. oscillatory protein dynamics, corresponding to a stable limit cycle in state space; and
iii. symmetric protein distributions, corresponding to a single stable fixed point in state space. The extent of these different regimes are indicated by the colored bars adjacent to each panel in Fig. 3.

**FIG. 5:**
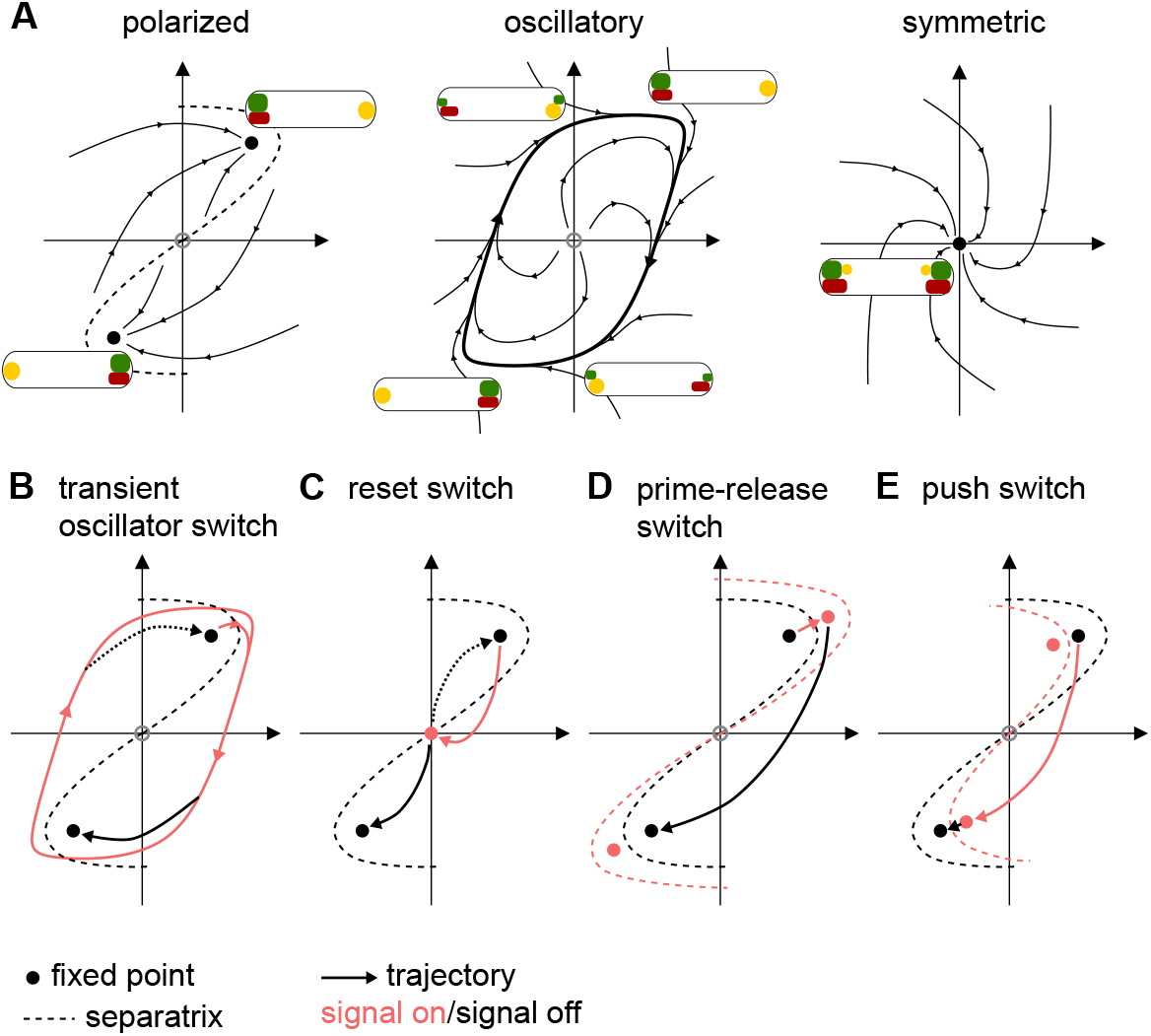
Schematic representation of the state space of the model and the dynamics during the switch. **A** The three qualitatively different state space topologies that are observed during the signal. The polarized state corresponds to a bistable system, the oscillatory state to a limit cycle and the symmetric state to a monostable system. **B-E** Changes in the state space induced by the switching signal by the mechanisms of the **B** transient oscillator switch, **C** reset switch, **D** prime-release switch and **E** push switch. Red lines indicate the state space and dynamics when the signal is turned on, black lines when the signal is switches off. In each case the system is initialized at the upper-right fixed point.

This analysis confirmed that band structures in Fig. 3 correspond largely to oscillatory dynamics in the presence of the *X* signal, while solid regions correspond to regimes where the system remains bistable when the signal is applied. However, we also identified regimes presenting two additional types of switches.

For large-amplitude signals, the system can transition from an oscillatory to a monostable regime. In this scenario, while the signal is applied the system gradually relaxes towards a symmetric configuration (Fig. 4C). Once the signal is removed, the system once again becomes polarized, but settles in the opposite polarization state from that in which it was initially. Effectively, the initial state of the system is erased and a new polarity state is chosen when the signal is removed. We therefore refer to this mechanism as a “reset” switch.

Finally, we found that as the oscillatory regime is approached, the onset of switching does not always coincide with the onset of oscillations. In the intervening region, the system still remains bistable. Examining the system trajectories, we observed qualitatively different behavior from Fig. 4B. Instead of switching once the signal is removed, the system begins to switch immediately when the signal is applied, and subsequently remains stably polarized in the opposite orientation (Fig. 4D). We refer to this mechanism as a “push” switch.

We have thus identified four distinct classes of switching dynamics, corresponding to four qualitatively different trajectories (Fig. 4). To understand these different mechanisms from a more general nonlinear dynamics perspective, we next ask how the topology of the phase space changes in each case. Prior to the application of the signal, the system is in a bistable configuration with two stable fixed points corresponding to the two possible polarity orientations (Fig. 5A). The subsequent behavior differs for each mechanism.

### Transient oscillator switch

In this class of switching, the system becomes oscillatory when the signal is applied, following the prescribed path of the limit cycle in state space. Upon removal of the signal, the phase space reverts to being bistable. The system will then relax to one of the polarized fixed points. Which fixed point is chosen depends on the state at the end of the signal period, and in particular on which side of the separatrix (the division between the basins of attraction of the two fixed points) the state lies, as illustrated in Fig. 5B. The duration of the signal relative to the oscillation period determines the phase at the time of signal removal and hence the final polarity state. How sensitive an oscillatory switch is to the signal duration varies dramatically between different regulations in our model, being relatively high for *k_b_* and *k_ab_*, but low for *k_bR_* and *k_r_* among others.

### Reset switch

Instead of following a limit cycle during the signal period, the reset switch gradually relaxes (usually along a spiraling trajectory) towards a single stable fixed point (Fig. 5C). Once again, the choice of polarity state upon removal of the signal depends only on which side of the separatrix the system is once the signal is removed. In the deterministic model, the choice of final polarity state is reliable even with a small remnant of asymmetry at the time of signal removal. However, this mechanism will be susceptible to noise in the protein dynamics that can overwhelm memory of the previous state (see below).

### Prime-release switch

This type of switch occurs when the model remains bistable even in the presence of the signal, and for parameter changes opposite to those that induce oscillations. The application of the signal does not cause a change in the topology of the state space, but does change the position of the fixed points and separatrix. If the signal is sufficiently strong, it may be that the new fixed points lie on the opposite side of the previous separatrix (Fig. 5D). However, since the current state remains on the same side of the new separatrix, the system simply relaxes to the new fixed point with the same polarity orientation (the “prime” phase). Only upon removal of the signal (the “release” phase) does the system find itself in the basin of attraction of the opposite polarity state.

This picture allows us to rationalize various observations about this switching mechanism. The amplitude of the signal must be sufficiently large that the new fixed point lies on the opposite side of the old separatrix, leading to a threshold in *X*_max_. The duration of the signal must be sufficiently long for the state of the system to move across the old separatrix, leading to a threshold in *τ*. Once these criteria are met, switching is insensitive to the signal amplitude and duration since the system can remain at the new polarized fixed point indefinitely.

### Push switch

The mechanism of the push switch is similar to that of the prime-release switch, but effectively with the order of events reversed. The application of the signal (“push”) again leads to a shift in the positions of the bistable fixed points and separatrix, but in the opposite direction (Fig. 5E). The system in its initial polarized state now finds itself on the opposite side of the new separatrix, from where it relaxes to the oppositely polarized fixed point. Upon removal of the signal, the system relaxes to the new slightly shifted fixed point but retains the same polarization. This mechanism is again largely robust to changes in the signal duration (after a threshold time needed for the initial relaxation phase), but occurs only for very small ranges of signal amplitudes in our model.

### Signals with slow edges

Both the prime-release and push switches described above rely on the fact that the signal appears and disappears very quickly, which causes a correspondingly fast change in the phase space. We expected that if the onset and removal of the signal were slower than the relaxation of the system, then the state of the system would be able to track the fixed points as they move gradually from their old to their new positions and no switching would occur. To test this prediction we computed the dynamics with the *X* signal increasing and decreasing gradually according to *X_t_*(*t*) = *X*_max_(1 − *e^−λt^*) for 0 ≤ *t* < *τ* and *X_t_*(*t*) = *X*_max_(1 − *e^−λτ^*)*e*^−*λ*(*t*−*τ*)^ for *t* ≥ *τ* (Fig. S3). We saw that for large *λ* ≫ 1, the dynamics was similar to a step signal and switching continued to occur (Fig. S4). However for slow signals with *λ* ≲ 1, switching in bistable regimes was abolished (Figs. S5 and S6). This was specific to the prime-release and push mechanisms since switching in oscillatory regimes continued to occur, with slight shifts to band boundaries reflecting the effects of the gradual signal on the oscillation phase (Figs. S7–S9).

### Stochastic effects

As seen in Fig. 3, the switching probability of the stochastic model for low to intermediate noise levels tends to closely follow the boundaries of regions in which the deterministic model switches (see also Figs. S10 and S11). However, switching can also occur for signal parameters *X*_max_ and *τ* for which the deterministic system does not switch. In particular, the regimes in which switching can occur are greatly expanded by noise for prime-release and push switches, while the transition boundaries between switching and non-switching regimes of relaxation oscillators appear much sharper. For reset switches, switching remains relatively robust with short signals, which are cut off before the system has fully relaxed to a symmetric state. For longer signals the switching probability approaches 0.5, as the new polarity state is chosen randomly once the signal is removed.

Fig. 6A shows the switching probability for the signal parameters indicated in Fig. 3 for increasing noise level. Each mechanism displays a different noise threshold at which the switching probability departs from the deterministic result (either 0 or 1 depending on the signal parameters). This threshold is highest for the transient oscillator (+, Δ), and lowest for the push (×) and prime-release switches (∗). Around *N* ≈ 10^3.5^ the switching probability converges to approximately 0.5 for all mechanisms. For higher noise levels (smaller *N*), all the mechanisms show qualitatively similar damped oscillations around 0.5. Similar behavior is observed in stochastic trajectories in the absence of any signal, indicating that these features are primarily the result of the dynamics during the period that the signal is not present (*τ* ≤ *t* ≤ *t*_end_). For this reason we first focus on the regime *N* ≳ 10^4^, in which the switching behavior remains influenced by the signal, and return to the high-noise behavior later.

**FIG. 6:**
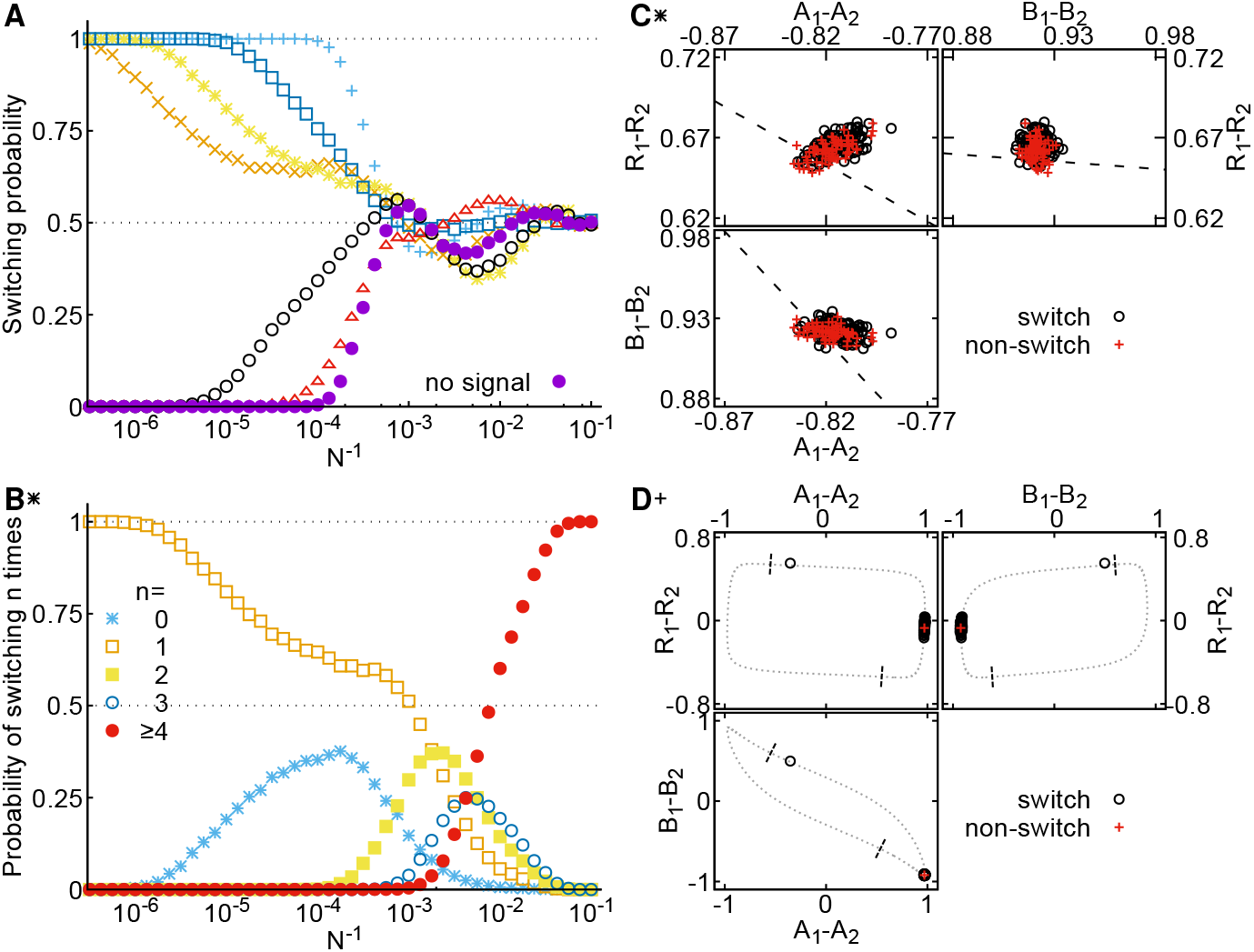
Behavior of the model at different noise levels. **A** Switching probability as a function of noise strength for different switching mechanisms. Signal parameters are indicated by the corresponding symbols in Figs. 3 and 4. Each data point represents the results of 10^4^ stochastic realizations. **B** Probability of different numbers of switching events at different noise levels for the prime-release switch. Signal parameters are as for Fig. 4B. **C** States of 200 stochastic realizations (*N* = 10^4^) of the prime-release switch at *t* = *τ*. Dashed line shows an estimate of the position of the separatrix in the absence of the signal (see Materials and Methods). **D** States of 200 realizations of the relaxation oscillator at *t* = *τ* (*N* = 10^4^). Dotted line shows the deterministic limit cycle of the system during the signal, dashed lines indicate where the limit cycle intersects with the separatrix in the absence of signal. The three different panels in **C** and **D** show different two-dimensional projections of the nine dimensional phase space. Point type and color indicate whether the system switches polarity (red) or not (black).

### Noise-induced switching errors

The switching probability, comparing only the states of the system before and after the signal is applied, cannot distinguish between cases in which noise prevents a switch from occurring and cases in which noise causes an extra switch to occur. We therefore examined the number of polarity switching events, defined as times at which *A*_1_ = *A*_2_, in stochastic trajectories. The distributions of such events are plotted for the prime-release switch in Fig. 6B (see Fig. S12 for the other cases). We observe that the initial decrease in switching probability around *N* ≈ 10^6^ corresponds to the appearance of a sub-population of realizations that do not switch. A flattening out of the switching probability around *N* = 10^4^, coincides with the appearance of trajectories exhibiting an extra second switch, due largely to stochastic switches during the period when no signal is present.

In the prime-release mechanism, switching is triggered by the removal of the signal. In the presence of noise, the system fluctuates around the fixed point of the dynamics, rather than resting exactly at the fixed point. The range of these fluctuations, visualized by sampling the states of different stochastic realizations at the end of the signal (prime) phase (Fig. 6C), expands with increasing noise strength. Importantly, when the signal is removed the states of the system are clustered close to the new separatrix of the system, allowing them to be forced from the basin of one fixed point to the other by noise. The same mechanism accounts for the expansion of the range of signals for which switching can be induced in the presence of noise beyond that in which the deterministic model will switch (Figs. 3 and 6A, ∘). The signal is not sufficiently strong for the deterministic fixed point with the signal applied to cross the original separatrix. However, some fraction of the distribution of states around this fixed point will lie close enough to the separatrix to undergo a switch when the signal is removed. Similar behavior can also be observed for the push signal with respect to the distribution of states at the onset of the signal period.

For the transient oscillator the initial deviation from the deterministic results is due to noise-induced extra switches once the signal has been removed. Noise in the dynamics during the signal predominantly leads to phase variability, as different realizations spread out around the limit cycle. However, the state of the system at the removal of the signal is typically far from the separatrix (Fig. 6D) in the slow phase of the dynamics where *R* reacts to the new polarity of *A* and *B*. Under these conditions, for the system to cross into the opposite basin of attraction requires extremely high noise levels. Hence, oscillatory switching appears extremely robust to noise.

### Coherence resonance

Returning to the high-noise regime (*N* ≲ 10^3.5^), where switching in the absence of any signal dominates, we observe that the switching probability oscillates before it saturates at 0.5 for very high noise (Fig. 6A). These oscillations are reminiscent of a so-called “coherence resonance” [24]. A coherence resonance occurs when the activation timescale for noise to drive the system across the separatrix of a bistable system becomes shorter than the relaxation timescale to reach the vicinity of the opposite fixed point. The trajectory of such a stochastic system has a largely oscillatory character. Indeed, the power spectrum of the dynamics changes from monotonically decreasing at small noise to peaked at a finite frequency for larger noise (Fig. 7A), indicating the appearance of oscillations. Additionally, the height of this peak shows a maximum at a finite noise level (Fig. 7B), confirming the coherence resonance behavior. Thus at high noise levels, noise can drive the system between the two polarity states with a largely oscillatory dynamics, even in the absence of any *X* signal (Fig S13).

**FIG. 7:**
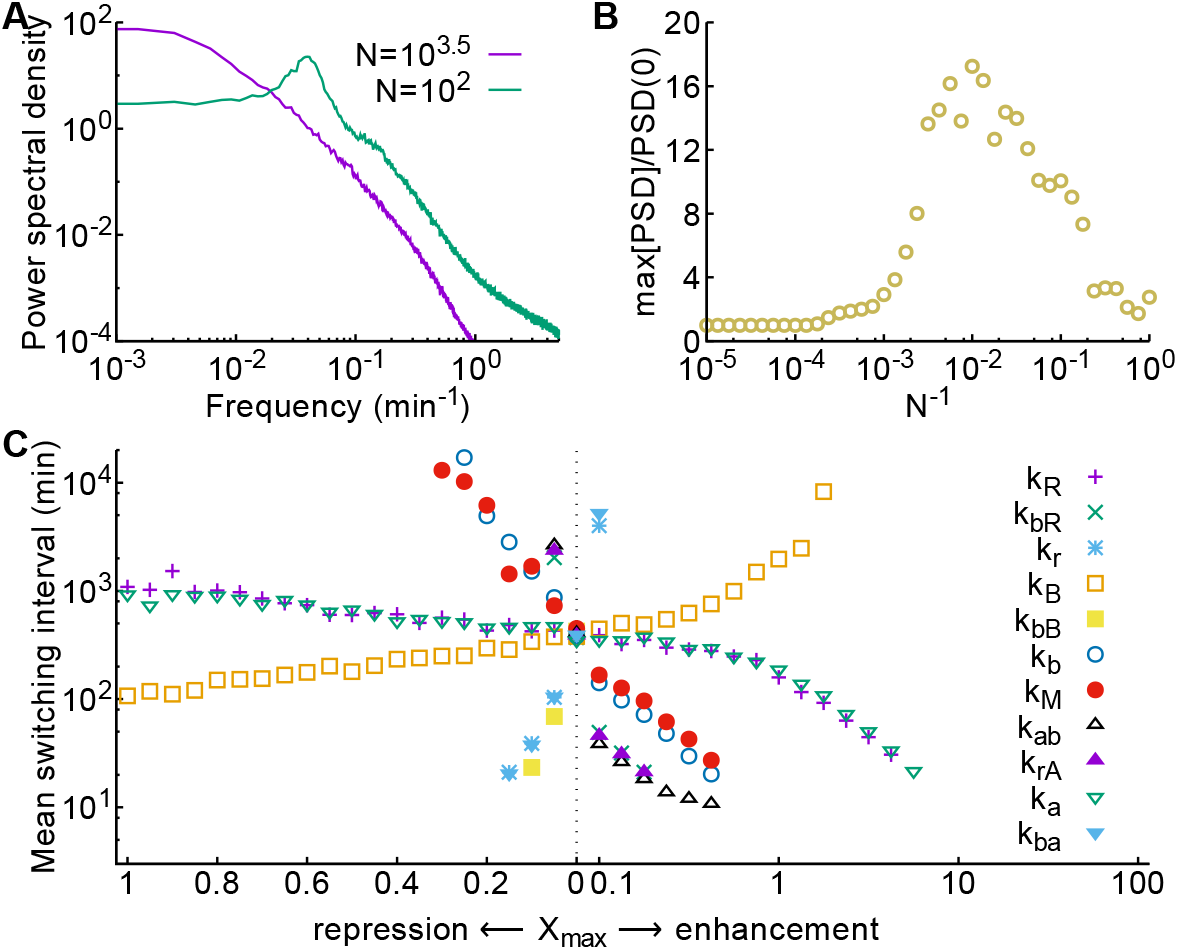
Stochastic switching. **A** Power spectral density of *A*_1_(*t*) − *A*_2_(*t*) in the absence of any *X* signal for different noise strengths. A peak in the power spectrum at high noise indicates stochastic coherence. **B** The maximal power density relative to the power at zero frequency shows a non-monotonic dependence on the noise strength. **C** The mean time between switching events, defined as points when *A*_1_ = *A*_2_, varies as different signals are applied, at a noise level *N* = 10^3.75^. Signals that generate deterministic oscillations have been excluded. Times between switches were extracted from stochastic trajectories with the signal applied continuously for 50000 min.

### Signal-induced stochastic switching

The application of a signal could also influence the stochastic switching rate during the period that the signal is active. For example, a signal could lower the height of the separatrix barrier between two fixed points, thereby increasing the chance of a stochastic switch. To study which signals could give rise to such an increase of stochastic switching, we analyzed long trajectories where signals with different amplitudes were applied continuously. Fig. 7C shows the resulting mean times between switching events. We observe that indeed the mean time between switches is affected by the choice of signal. Interestingly, the mean interval between switches decreases as the signal approaches the regime of oscillations, consistent with a reduction in the height of the separatrix barrier as the bifurcation point is approached. Conversely, switches become rarer when the signal varies in the opposite direction, into the prime-release regime. In general, however, the frequency of switching is extremely low such that the expected number of switches during one signal period approaches zero. We therefore conclude that the effects of stochastic switching during the signal will be negligible and dominated by the responses of the system to the transient phases of the signal.

## DISCUSSION

In this work we have developed a general classification of signal-induced polarity switching mechanisms based only on how a signaling protein changes the topology of a polarity system’s state space. This classification consists of four qualitatively different switching mechanisms: (i) the transient oscillator switch, (ii) the reset switch, (iii) the prime-release switch, and (iv) the push switch. From a nonlinear dynamics perspective, the four types of behavior in Fig. 5 appear to exhaust the spectrum of possible behaviors, such that we do not expect additional classes to appear in other polarity models. Indeed, it is somewhat surprising that the interaction scheme of the Guzzo et al [21] model for *M. xanthus* polarity, which we took as the starting point for our analysis, is capable of producing all four types of switches. It remains to be seen whether the capacity for such diverse switching phenomonology is common to other models of prokaryotic and eukaryotic cell polarity, and which features of such models enable different switching modes. Some models, in particular those with fewer components, are likely not able to produce all four types of switches, e.g. because they cannot generate oscillations.

We showed how the different mechanisms respond to signal variability and internal molecular noise. For instance, while the transient oscillator switch is most sensitive to signal variability it is least sensitive to molecular noise. By contrast, the prime-release switch is least sensitive to signal variability, but very sensitive to molecular noise. These differences in behavior will be useful as signatures to identify the actual switching mechanisms in biological systems. In addition, these properties will be relevant for the design of synthetic systems.

One of the main predictions of our analysis is that the cells’ behavior during a long signal is characteristic for the different switching mechanisms. In systems where the signal can be either monitored or externally controlled, the mechanism of the switch can be revealed by studying the relative timing between the signal and the reversal behavior of cells for which the signal is always present. For the prime-release switch, the polarity reversal can only occur after the signal is removed. The reset-switch has a characteristic loss of polarity during the reversal. If the reversal is observed while the signal is still present, the prime-release switch can be excluded. However, if the reversal occurs after the signal is removed, we cannot exclude the transient oscillator and the push switch, as it is possible that the reversal is triggered during the signal but happens only after the signal is removed. Note that such experiments will only give insight on the type of switch and not on the detailed interaction between the signaling protein and the polarity proteins, since the same qualitative dynamics can be generated by different modes of action of the signal.

Our analysis of the systems’ dynamics also revealed that, while the timing with respect to the input signal is different for the different mechanisms, the order in which the proteins of the core polarity system switch poles is almost always the same. This indicates that we cannot infer the interaction of the signaling protein *X* from looking at the order in which the polarity proteins switch poles, but that the order of switching is rather a characteristic of the interactions between different polarity proteins.

Finally, by analogy with the paradigmatic genetic toggle switch [25], the functionality analyzed here can be regarded as a ‘spatial toggle switch’. The core of the genetic toggle switch is a circuit of two mutually repressing genes, conceptually similar to the mutual inhibition between the polarity marker *A* and its antagonist *B*. Some of the behavior is also analogous, e.g. molecular noise can cause the genetic toggle switch to flip spontaneously [26], just as it does for the polarity system. However, while the genetic toggle switch is a well-mixed bistable system, the core polarity system is a spatially extended bistable system that forms asymmetric patterns. The spatial extension of the polarity system allows a global signal (*X_t_*) to be converted into a local signal (differential activity of *X* at the two poles), in a way that would be impossible in a well-mixed system. This permits the polarity system to function as a true toggle switch, i.e. the same signal causes switching in both directions, in contrast to the original genetic toggle switch, where different signals “set” and “reset” the switch [25]. The true toggle (or “push-on push-off”) functionality in genetic switches requires more elaborate regulatory circuitry that manipulates the bistable system as a function of input signals to achieve control of the system [27–30]. By comparison, the control of pattern forming systems is only beginning to be explored, opening interesting directions for future research.

## METHODS

### Deterministic dynamics

Reaction rates *k_j_* were chosen as in [21], with the rate *k_ab_* = 15 min^−1^. The deterministic dynamics was computed with Mathematica (Wolfram Research Inc.) using the function NDSolve separately in each domain (before, during and after the signal), with initial conditions set according to the protein abundances at the end of the previous segment.

### Stochastic model

For the stochastic version of the model we used a Langevin extension of Eqs. 2, adding a noise term to each equation,

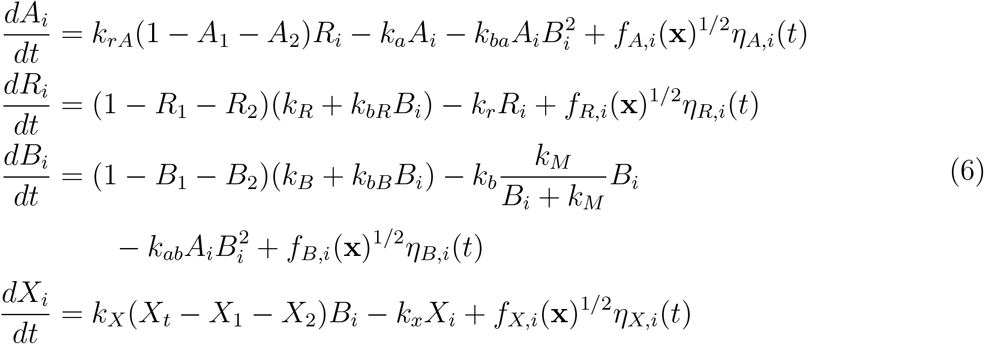

where **x** = (*A*_1_, *A*_2_, *R*_1_,…, *X*_2_) is the state vector, and the *η*_·,*i*_ are independent Gaussian random variables, 〈*η*_·,*i*_(*t*)〉 = 0 and 〈*η_p,i_*(*t*)*η_q,j_*(*t*′)〉 = *N*^−1^*δ_p,q_δ_i,j_δ*(*t* − *t*′). We have introduced *N* as a parameter to tune the magnitude of the noise, with the deterministic model being recovered as *N* → ∞. We chose to make the noise multiplicative by having the strengths *f*_·,*i*_(**x**) depend on the current state of the system, **x**. Specifically,

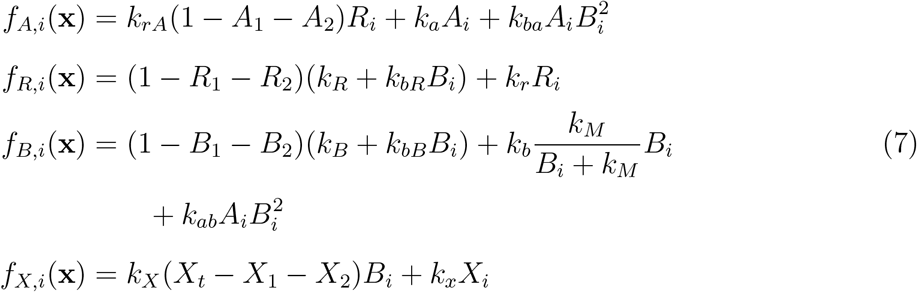

We emphasize here that these noise terms were chosen simply as one plausible generalization of Eqs. 2. While they resemble those that might be obtained from a system-size expansion of a full Master equation for the reactions underlying Eqs. 2 [31, 32], we note that since the original model is defined only in terms of the rate equations and not in terms of the underlying molecular reactions, no such systematic derivation of the noise is possible. We verified that the particular choice of the form of the noise did not affect our conclusions, and found qualitatively similar results when white noise was used (implemented by fixing **x**= (1/3, 1/3,…, 1/3, *X_t_*/3, *X_t_*/3) in Eq. 7), see Figs. S14 and S15.

Simulations of the stochastic model were performed by directly integrating Eqs. 6 using an update rule of the form

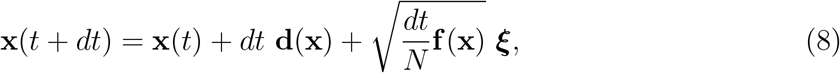

where **d**(**x**) represents the deterministic part of Eqs. 6, **f**(**x**) = (*f_A,_*_1_, *f_A,_*_2_, *f_R,_*_1_,…, *f_X,_*_2_) is a vector of noise strengths, ***ξ*** is a vector of independent samples from a normal distribution, and multiplication of **f**^1/2^ and ***ξ*** is performed elementwise. A time step *dt* = 10^−4^min was used throughout. After each update step, all protein abundances were corrected such that none were negative or exceeded the total protein numbers (i.e. ensuring *A*_1_ + *A*_2_ ≤ 1, and similarly for each other protein). The simulation code (implemented in C++) is available at github.com/gerland-group/langevin switching.

### Estimation of separatrices

The separatrix lines in Fig. 6C,D were estimated as follows. For the prime-release switch (Fig. 6C), we first estimated the state space around the fixed point in the presence of the signal by simulating 10000 stochastic trajectories with *N* = 10^3^ until *t* = *τ*. For each of these points, we determined on which side they fell of the separatrix in the absence of signal, by taking these as the initial conditions for deterministic simulations over the period *τ* ≤ *t* < *t*_end_. The projections of the separatrix in the planes shown was then estimated by using a linear discriminant classifier to determine, for each of the two-dimensional projections of the data in turn, the decision boundary between the sets of states that belong to each of the basins of attraction. This analysis was performed using the LinearDiscriminantAnalysis class from scikit-learn [33] with default parameters. For the relaxation oscillator (Fig. 6D), we identified the path of the limit cycle from the trajectory of the deterministic model. The intersection points with the separatrix were then estimated by initializing simulations with the signal removed at different points along the limit cycle.

### Power spectra

Power spectral densities were estimated from trajectories sampled every 0.01 min for 50000 min by Welch’s method of averaged periodograms from overlapping segments of the trajectory [34] using the MATLAB (Mathworks) function pwelch with segments of length 2^16^ samples.

## Acknowledgments

We thank Tam Mignot and Sean Murray for helpful discussions. This work was supported by the German Research Council (DFG) within the framework of the Transregio 174 “Spatiotemporal dynamics of bacterial cells” (to L.S.-A. and U.G.) and within the framework of the Grauduate school for Quantitative Biosciences Munich (QBM) (to M.W.), by the Volkswagen Foundation (to U.G.), by the Max Planck Society (to L.S.-A.), and by the Joachim Herz Foundation (to M.W.).

## Author contributions

All authors designed research. F.T. and M.W. performed simulations and analyzed data. F.T., M.W. and U.G. wrote the paper.

