## Supplementary Figures for "Four different mechanisms for switching cell polarity"

### Supplementary Information:
### Four different mechanisms for switching cell polarity

Filipe Tostevin,[1,*] Manon Wigbers,[1,2,*] Lotte Søgaard-Andersen,[3] and Ulrich Gerland[1,†]

[1]*Physics of Complex Biosystems, Physics Department,*
*Technical University of Munich, James-Franck-Str. 1, 85748 Garching, Germany*

[2]*Arnold Sommerfeld Center for Theoretical Physics and Center for Nanoscience,*
*Ludwig-Maximilians-Universität München,*
*Theresienstr. 37, 80333 München, Germany*

[3]*Max Planck Institute for Terrestrial Microbiology,*
*Karl-von-Frisch-Str. 10, 35043 Marburg, Germany*

Keywords: cell polarity | signal transduction | toggle switch

[*]Contributed equally to this work.

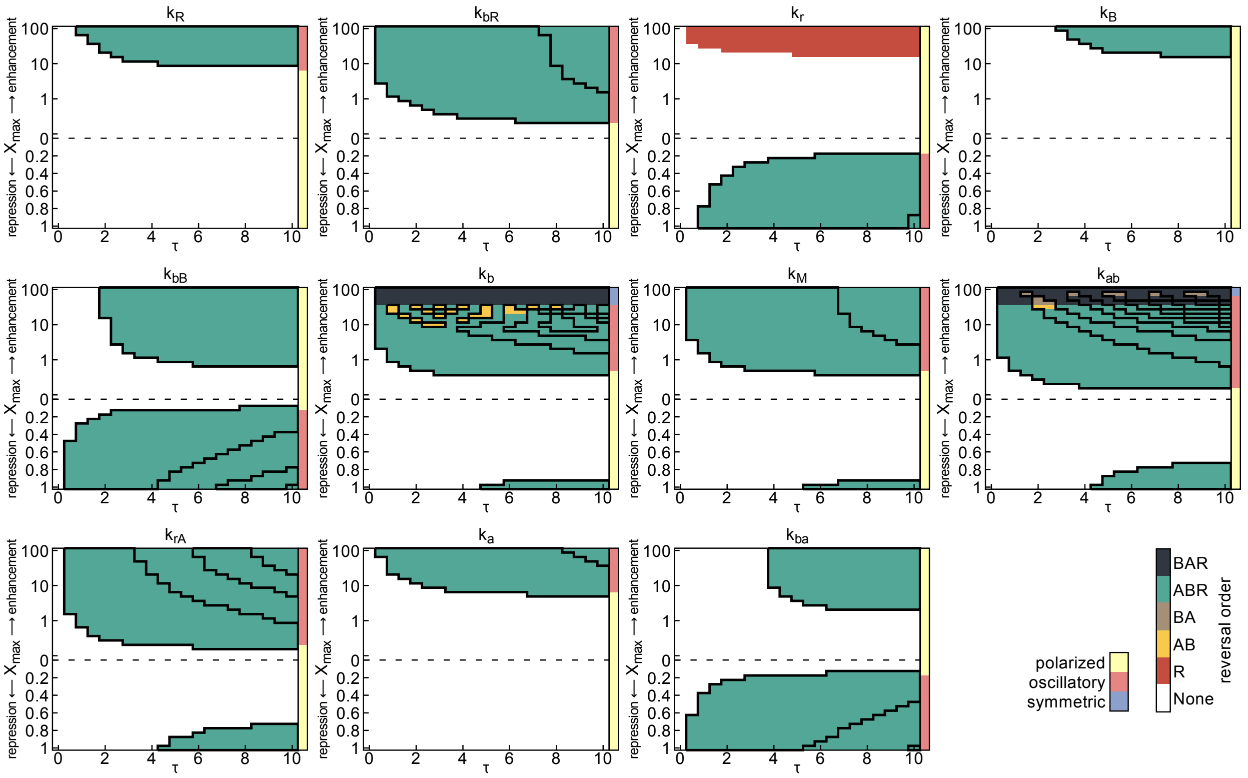


FIG. S1: Order of switching. Switching trajectories are obtained from the deterministic model. Black solid lines in the phase diagrams show switching regimes as in Fig. 3. The colors indicate in which order $A$, $B$ and $R$ switch polarity. In the regimes where the system switches polarity multiple times (due to the transient oscillator switch), the switching order represents the order of the first switch.

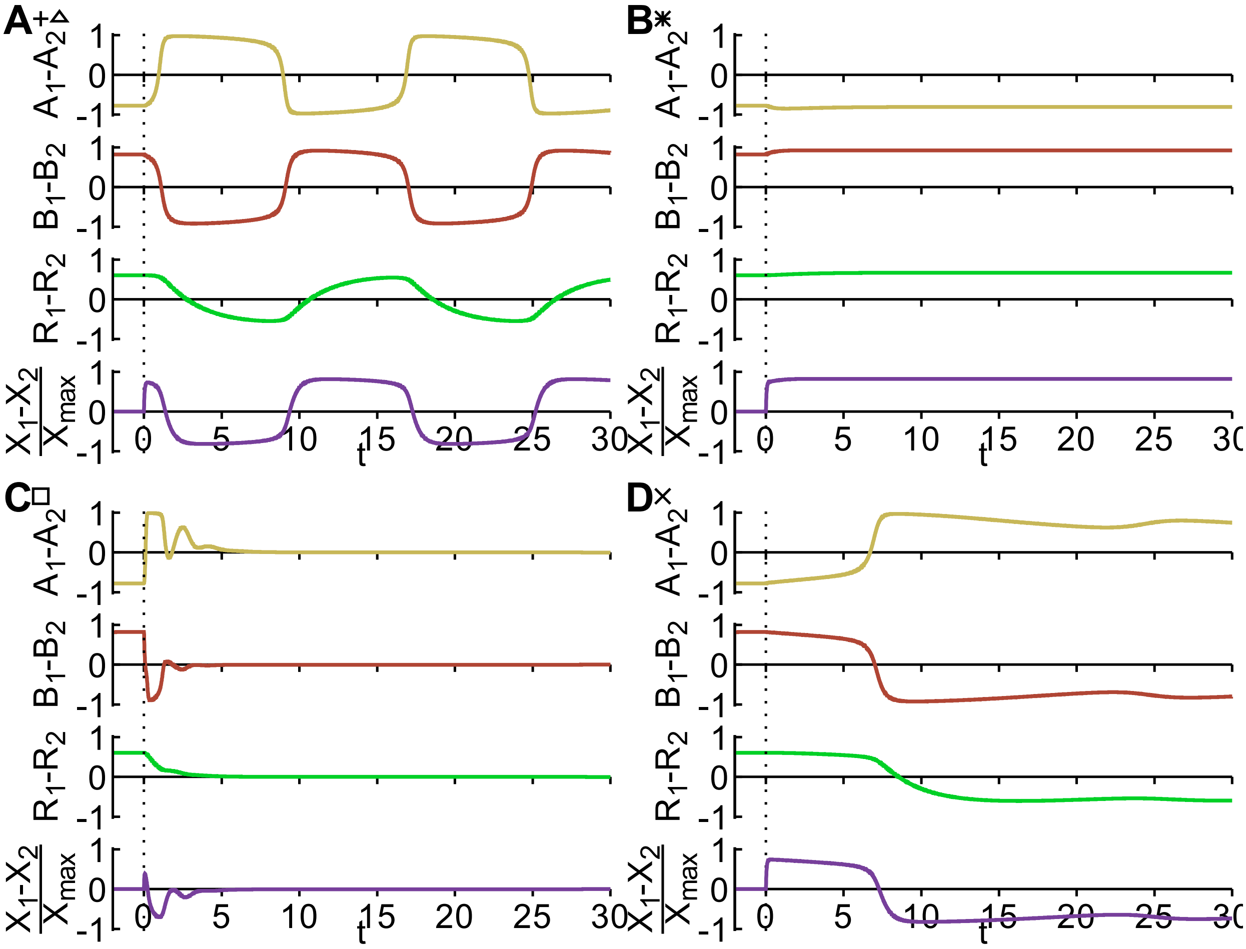


FIG. S2: Trajectories of the system during the signal for **A** the transient oscillator switch, **B** the prime-release switch, **C** the Reset switch and **D** the push switch. The symbols next to the panel labels indicate the signal parameter $X_{\max}$ as indicated in Fig. 3. The signal is applied for the duration of the simulation. During the transient oscillator switch (**A**) the polarity of the system oscillates, while during the reset switch (**C**) there is no polarity, i.e. the distribution of the proteins at pole 1 and pole 2 is symmetric. During the prime-release and push switch (**B** and **D**) the system is polarized during the switch.

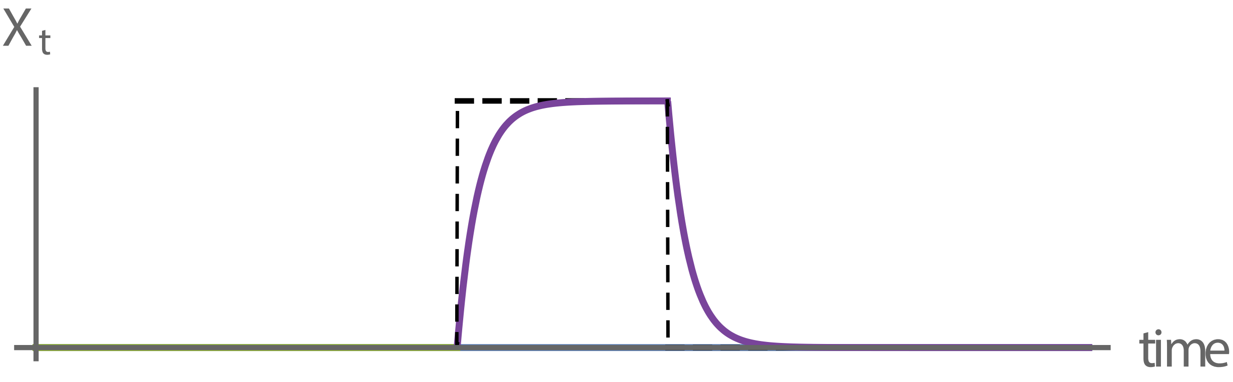


FIG. S3: Example of a gradually increasing and decreasing signal. The total amount of $X$, $X_t$, increases according to $X_t(t) = X_{\max}(1 - e^{-\lambda t})$ for $0 < t < \tau$ and decreases according to $X_t(t) = X_{\max}(1 - e^{-\lambda\tau})e^{-\lambda(t-\tau)}$ for $t > \tau$. The dashed line indicates the step-like signal.

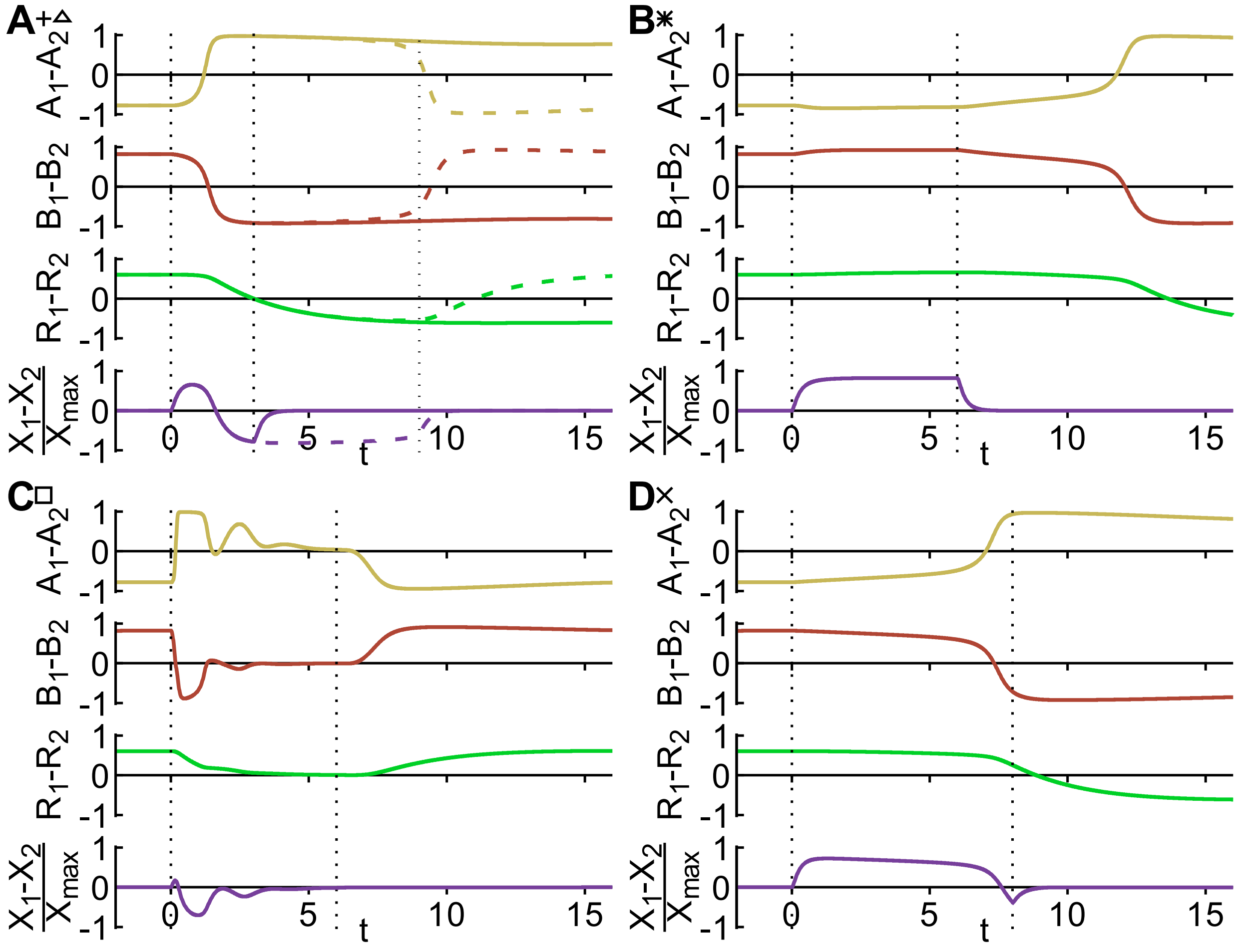


FIG. S4: Trajectories for a gradually increasing and decreasing signal with $\lambda = 4$. Signal amplitude $X_{\mathrm{max}}$ and duration $\tau$ are chosen the same as in Fig. 4, where in **A** the solid line corresponds to the short signal (plus-symbol) and the dashed line to the long signal (open triangle). The system shows qualitatively the same behavior as for the step-like signal.

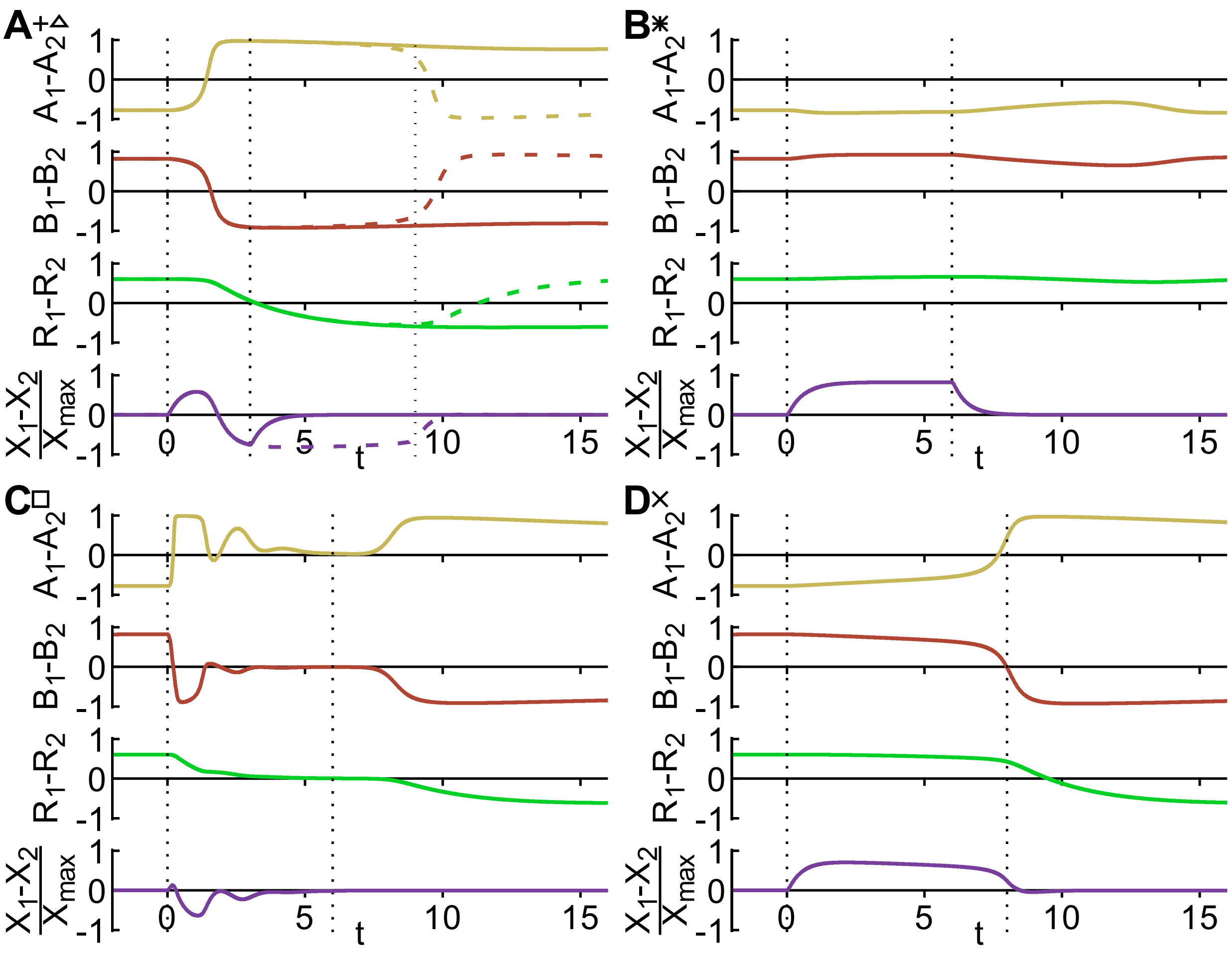


FIG. S5: Trajectories for a gradually increasing and decreasing signal with $\lambda = 2$. Signal amplitude $X_{\max}$ and duration $\tau$ are chosen the same as in Fig. 4, where in **A** the solid line corresponds to the short signal (plus-symbol) and the dashed line to the long signal (open triangle). For these gradual signals, the transient oscillator switch (**A**), the reset switch (**C**) and the push switch (**D**) switch qualitatively the same as for a step-like signal, while the prime-release switch (**B**) does not respond to the gradual signal.

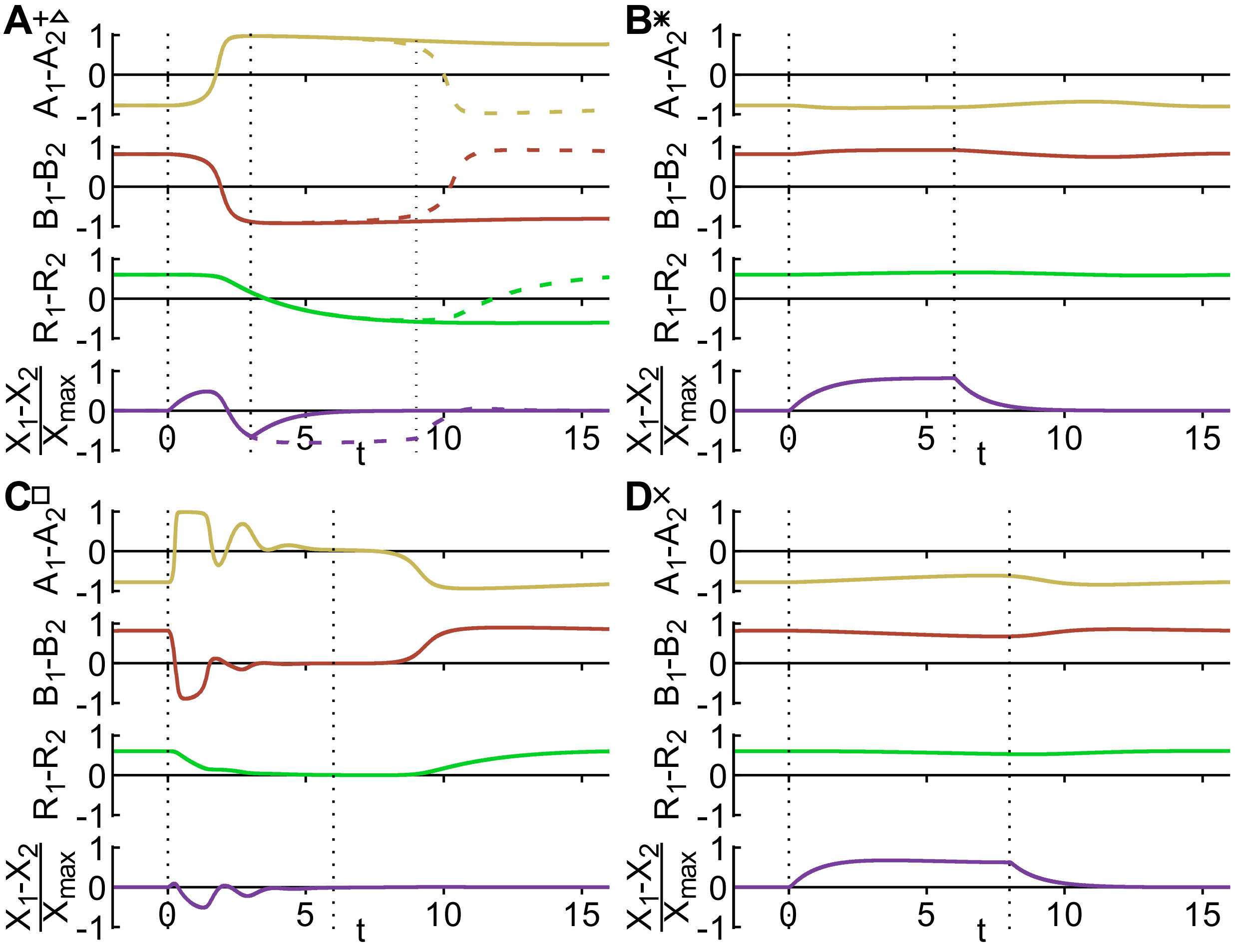


FIG. S6: Trajectories for a gradually increasing and decreasing signal with $\lambda = 1$. For these gradual signals, the prime-release (**B**) and push switch (**D**) do not respond to the signal, while the transient oscillator (**A**) and reset switch (**C**) do.

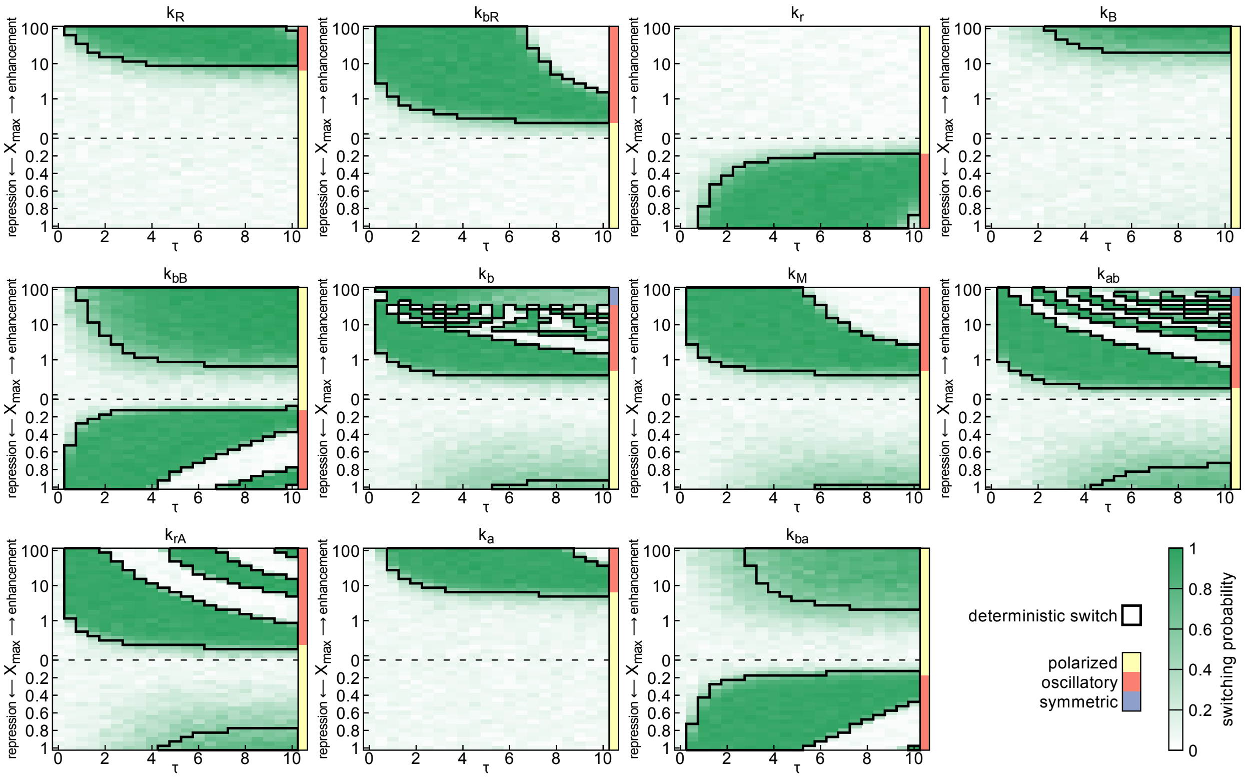


FIG. S7: Switching regimes for a gradually increasing and decreasing signal with $\lambda = 4$. Regions in which the deterministic model shows switches are indicated by thick black outlines. The green shading shows the switching probability of the stochastic model with $N = 10^3.75$. The upper half of the phase diagram shows results for a signal that enhances the reaction rate, and the lower half for a repression of the rate. The colored bars to the right of each panel indicate the class of dynamics when the corresponding amplitude of signal is applied, with yellow for polarized, orange for oscillatory and blue for symmetric polar distribution of $A$, for a gradually increasing and decreasing signal. The switching regimes are similar to the regimes for a step-like signal as shown in Fig. 3.

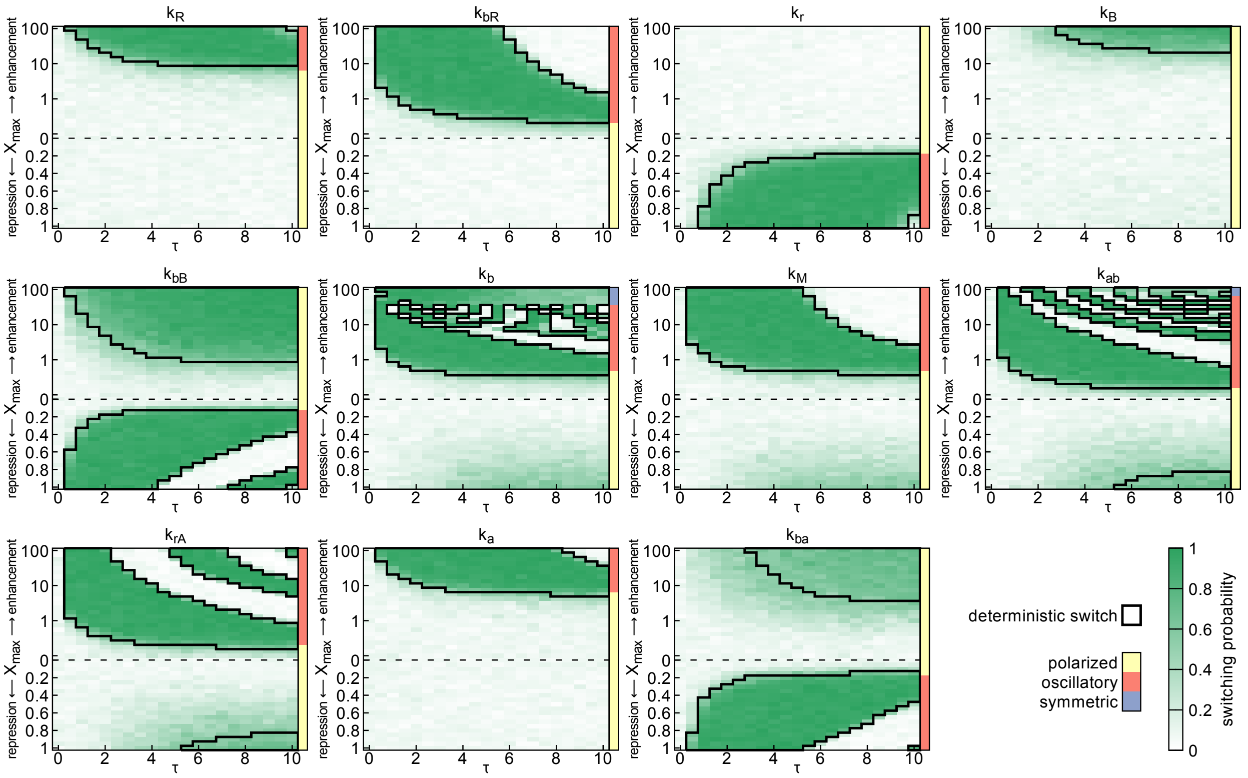


FIG. S8: Switching regimes for each of the model parameters for a gradually increasing and decreasing signal with $\lambda = 2$. The regimes where the prime-release switch acts to switch the polarity, for example via repression of the parameter $k_{ab}$ or $k_{rA}$, have become smaller.

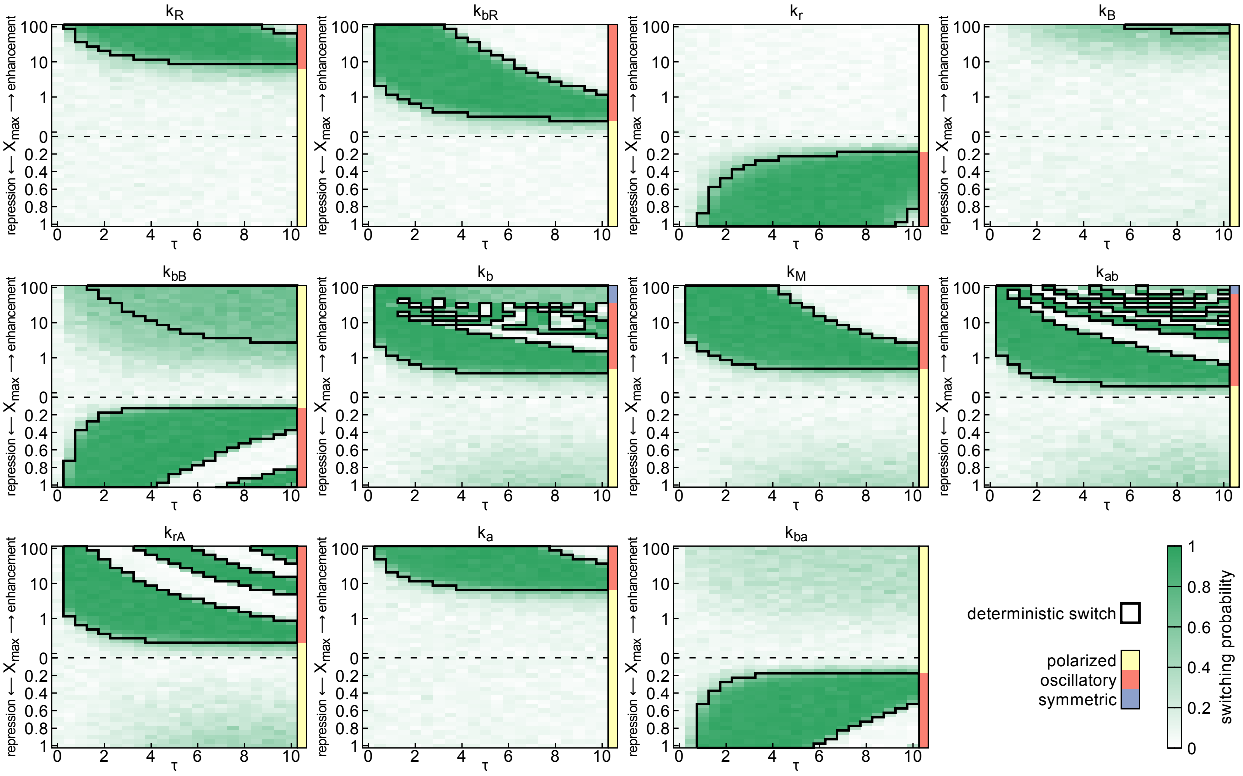


FIG. S9: Switching regimes for each of the model parameters for a gradually increasing and decreasing signal with $\lambda = 1$. The regimes where the prime-release switch acts to switch the polarity becomes smaller, for example by enhancing $k_B$, or completely vanishes, for example via repression of the parameters $k_{\rm ab}$ or $k_{\rm rA}$. In addition, the regimes where the push switch acts vanishes, for example via a slight enhancement of the parameter $k_{\rm rA}$.

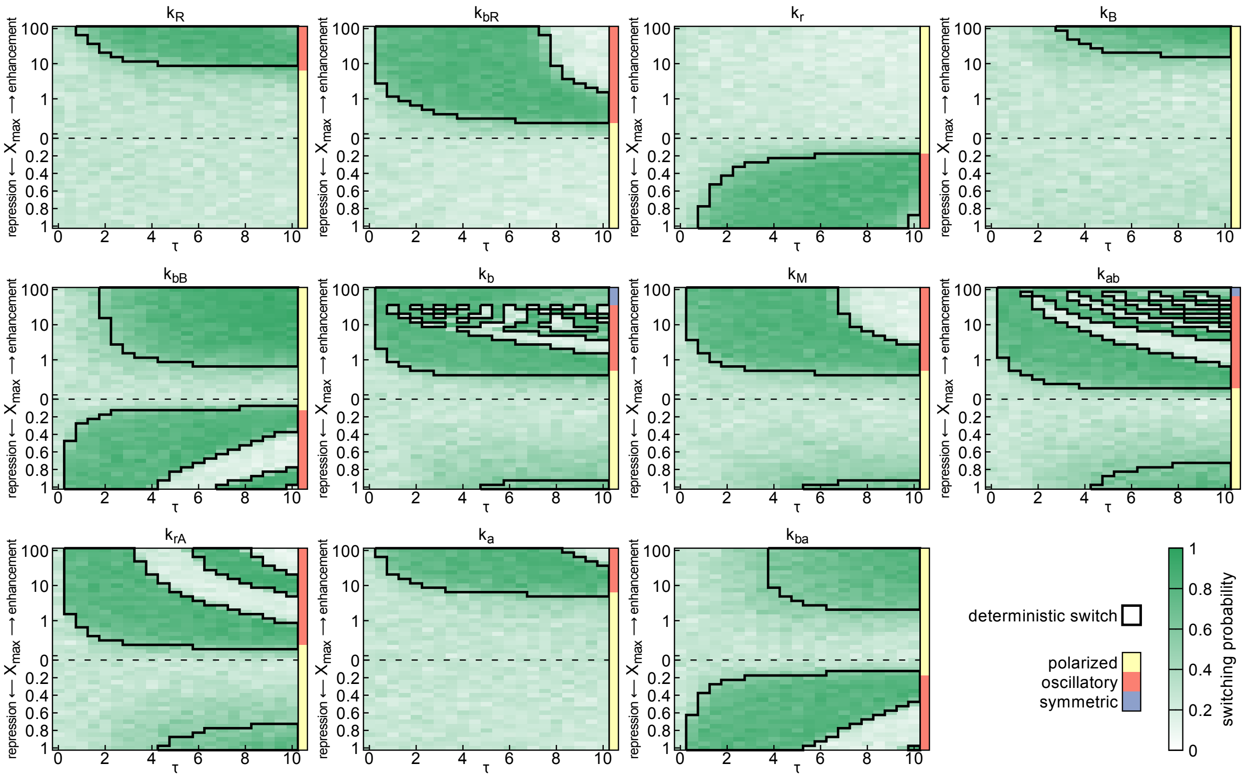


FIG. S10: Switching regimes for each of the model parameters with a step-like increasing and decreasing signal. The green shading shows the switching probability of the stochastic model with $N = 10^{3.5}$. The stochastic switching probability, outside of the deterministic switching regimes (solid black lines), is higher as compared to a noise level of $N = 10^{3.75}$ as shown in Fig. 3, while the switching probability in the deterministic regimes is smaller.

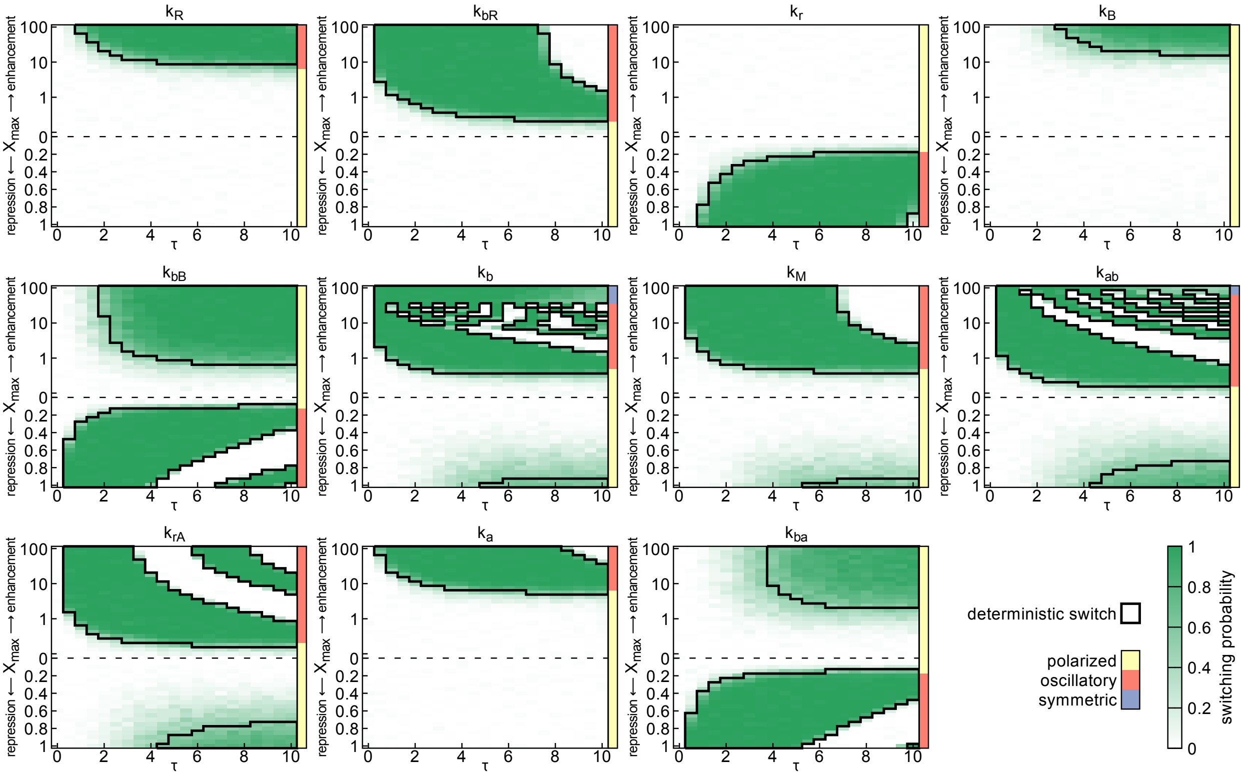


FIG. S11: Switching regimes for each of the model parameters with a step-like increasing and decreasing signal. The green shading shows the switching probability of the stochastic model with $N = 10^4$. The stochastic switching probability, outside of the deterministic switching regimes (solid black lines), is smaller as compared to a noise level of $N = 10^{3.75}$ as shown in Fig. 3, while the switching probability in the deterministic regimes is higher.

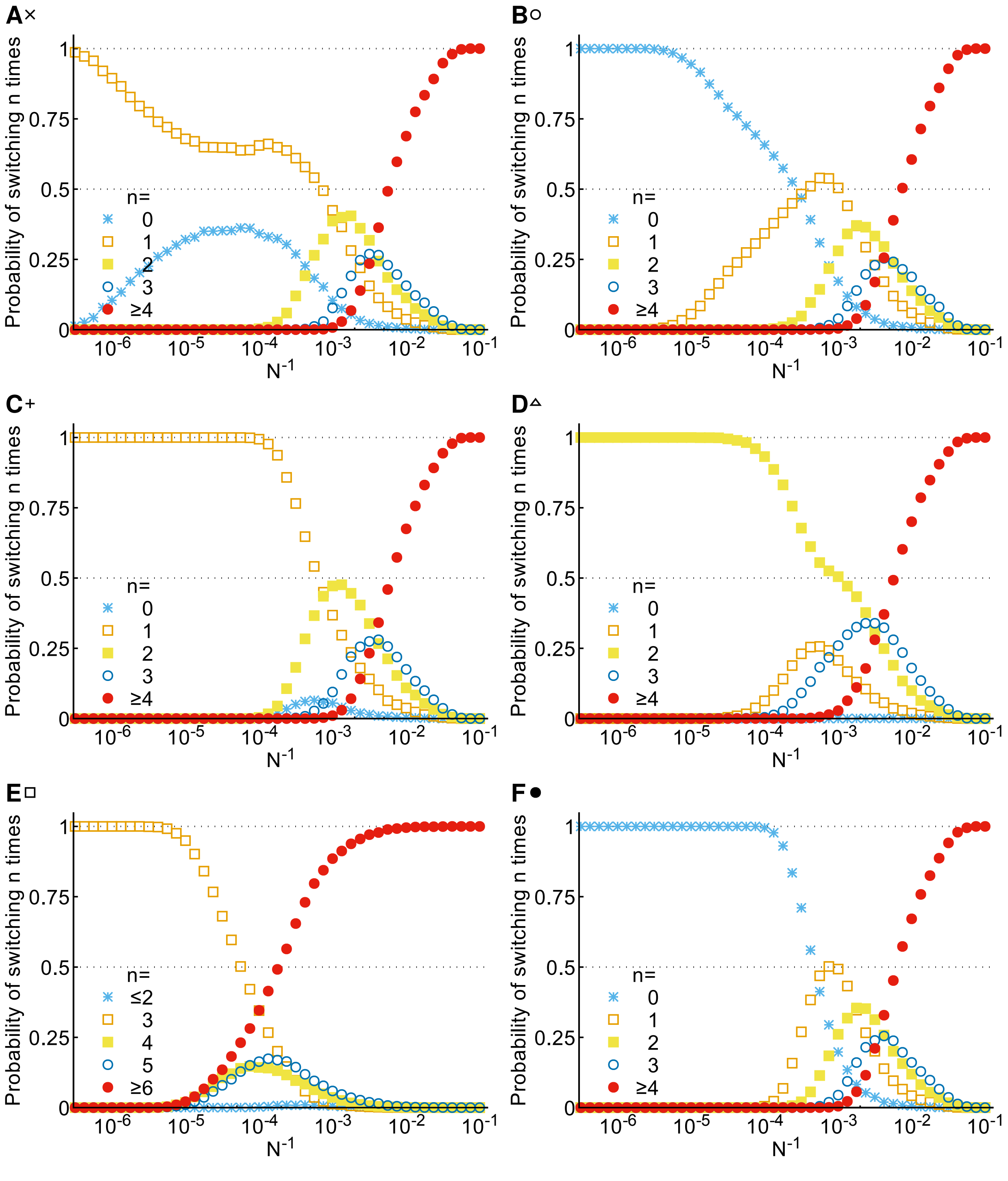


FIG. S12: Probability of different numbers of switching for different noise levels. Symbols next to the panel labels **A-E** correspond to the signal amplitude and duration as shown in Fig. 3. **F** shown the probability of different numbers of switches without a signal.

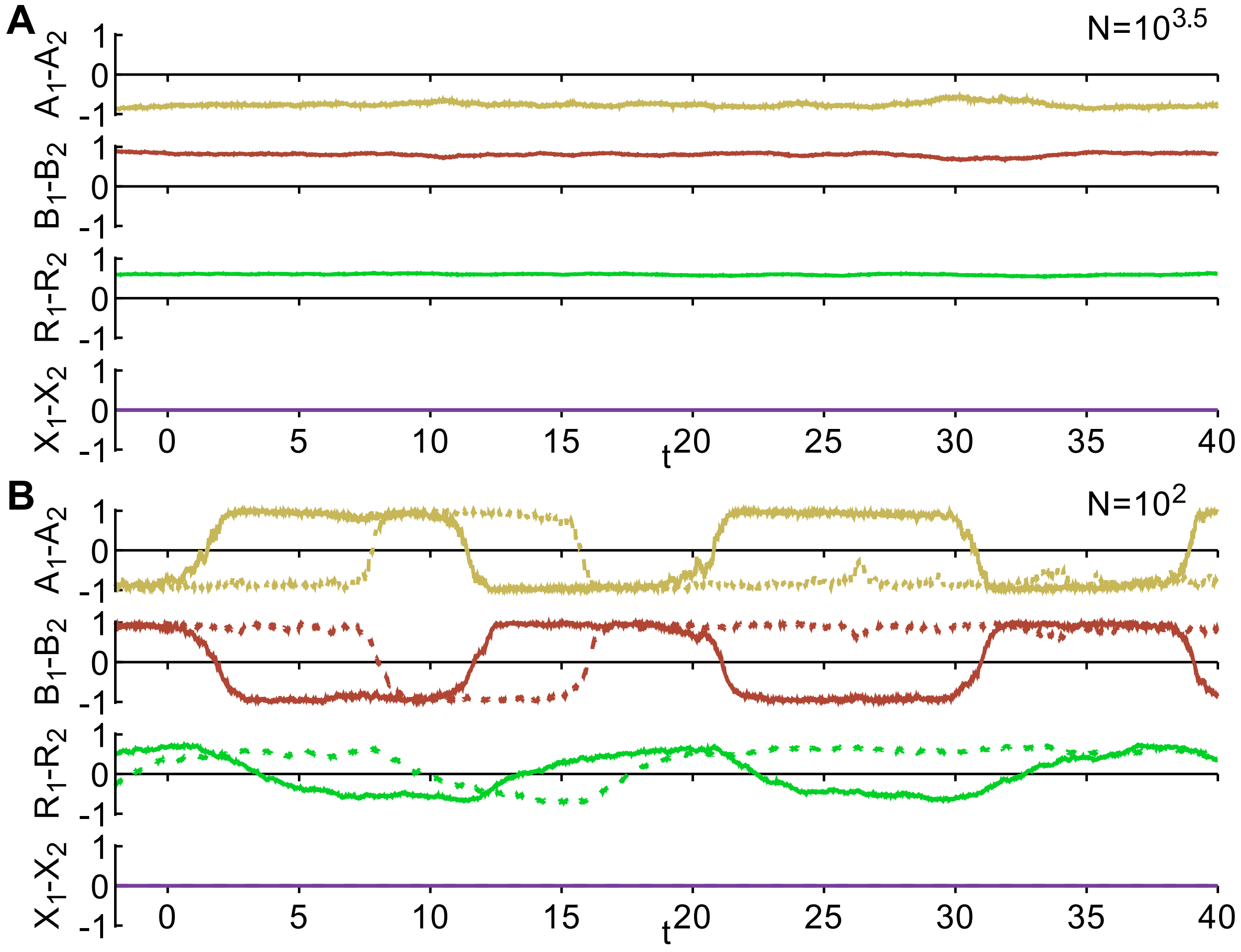


FIG. S13: Polarity switching of the stochastic model without a signal. **A** for low noise levels ($N = 10^{3.5}$) the system does not switch for the duration of the simulation. **B** for high noise levels ($N = 10^2$) the polarity switches several times without applying a signal.

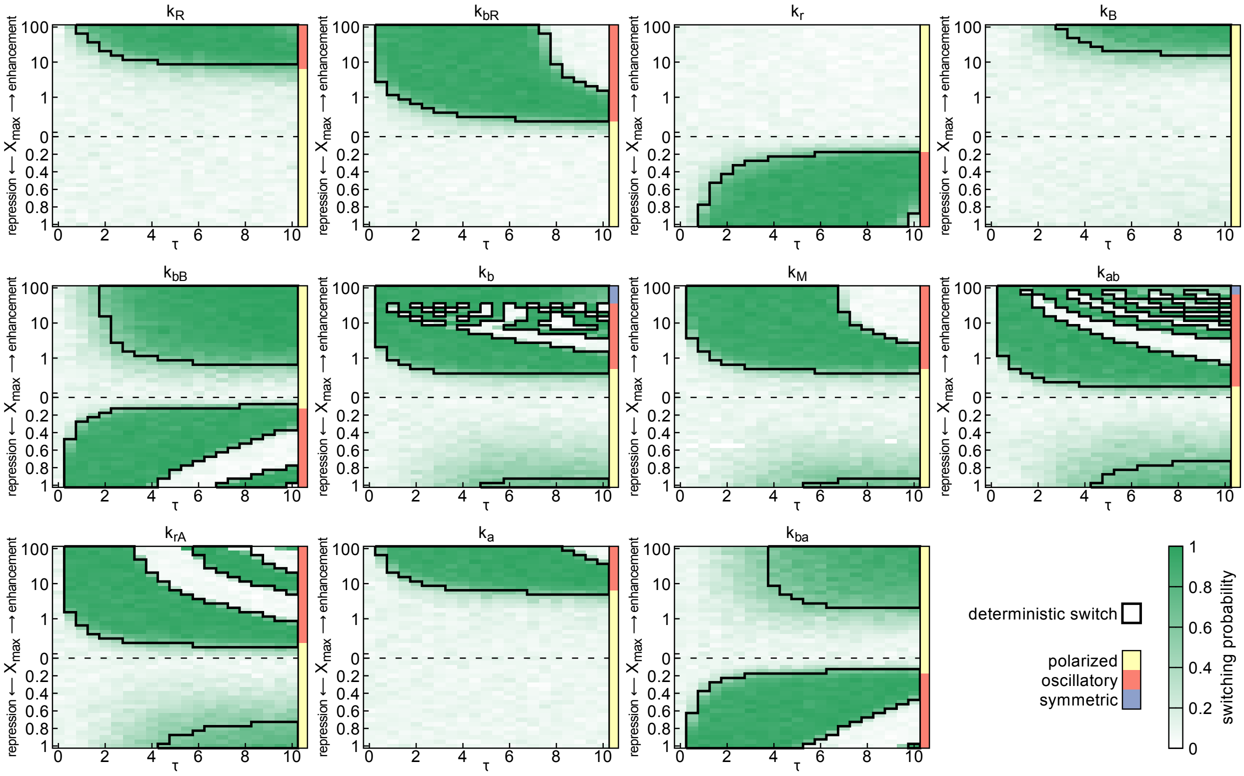


FIG. S14: Switching regimes for each of the model parameters with a step-like increasing and decreasing signal. The green shading shows the switching probability of the stochastic model with white noise and with $N = 10^4$. The switching regimes are qualitatively similar to the switching regimes in Fig. 3.

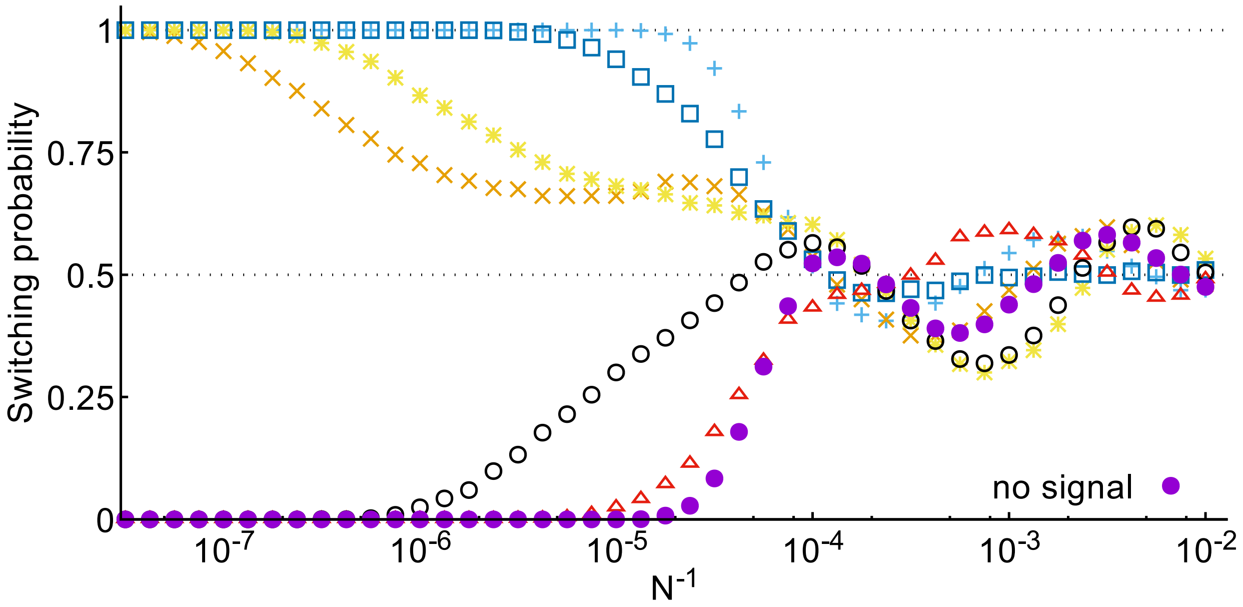


FIG. S15: Switching probability of the stochastic model with white noise. The signal parameters are indicated by the corresponding symbols in Figs. 3 and 4. Results are qualitatively similar to the results presented in Fig. 6A.
